# Heterophilic interactions between cell-surface proteins Teneurin-m and Capricious drive dendrite segregation in the *Drosophila* olfactory circuit

**DOI:** 10.64898/2026.04.22.719985

**Authors:** Hui Ji, Jingxian Li, Yizhen Xu, Kenneth Kin Lam Wong, Yunming Wu, David J. Luginbuhl, Yanbo Zhang, Zhuoran Li, Jaeyoon Lee, Robert C Jones, Stephen R Quake, Demet Araç, Engin Özkan, Liqun Luo

**Affiliations:** Department of Biology, Stanford University, Stanford, CA 94305, USA; Howard Hughes Medical Institute, Stanford University, Stanford, CA 94305, USA; Department of Biochemistry and Molecular Biology, The University of Chicago, Chicago, IL 60637, USA; Neuroscience Institute, The University of Chicago, Chicago, IL 60637, USA; Institute for Biophysical Dynamics, The University of Chicago, Chicago, IL 60637, USA; Center for Mechanical Excitability, The University of Chicago, Chicago, IL 60637, USA; Biology Graduate Program, Stanford University, Stanford, CA 94305, USA; Department of Bioengineering, Stanford University, Stanford, CA 94305, USA; Department of Applied Physics, Stanford University, Stanford, CA 94305, USA; The Chan Zuckerberg Initiative, Redwood City, CA 94063, USA

## Abstract

How dendrites of different neurons segregate into discrete spatial domains during neural circuit assembly is poorly understood. Here, using the *Drosophila* olfactory system, we found that heterophilic interactions between two cell-surface proteins Teneurin-m (Ten-m) and Capricious (Caps) drive dendrite segregation. Ten-m and Caps are expressed in largely inverse patterns across projection neuron (PN) types when PNs are establishing their dendritic territories. Loss of Ten-m in Ten-m^+^ PNs causes their dendrites to invade Caps^+^ territories, whereas loss of Caps in Caps^+^ PNs causes dendrite invasion into Ten-m^+^ territories. Structure-guided mutations that abolish Ten- m–Caps binding disrupt dendrite segregation, whereas the same mutation on Ten-m preserves its homophilic attraction in a synaptic partner matching assay. These results support a model in which mutual repulsions between two inversely expressed cell-surface proteins drive dendrite segregation into discrete glomerular territories.

## Introduction

The assembly of functional neural circuits requires neurons to elaborate dendritic arbors that occupy specific territories. In many systems, dendrites of different neuronal types segregate into discrete, non-overlapping spatial domains^1,2^. Whereas mechanisms governing self-avoidance—the segregation of sister branches within a single neuron—are well characterized^3,4^, the molecular logic driving the segregation of dendrites from different cell types remains far less understood.

The *Drosophila* olfactory system offers a tractable model to study dendrite segregation. In the antennal lobe, ∼50 types of olfactory receptor neurons (ORNs) form one-to-one synaptic connections with ∼50 types of projection neurons (PNs) at ∼50 discrete glomeruli^5^. PN dendrites pre-pattern the antennal lobe before ORN axons arrive^6–8^, suggesting that dendrite–dendrite interactions play a critical early role in establishing the glomerular map (Figure 1A).

**Figure 1.**
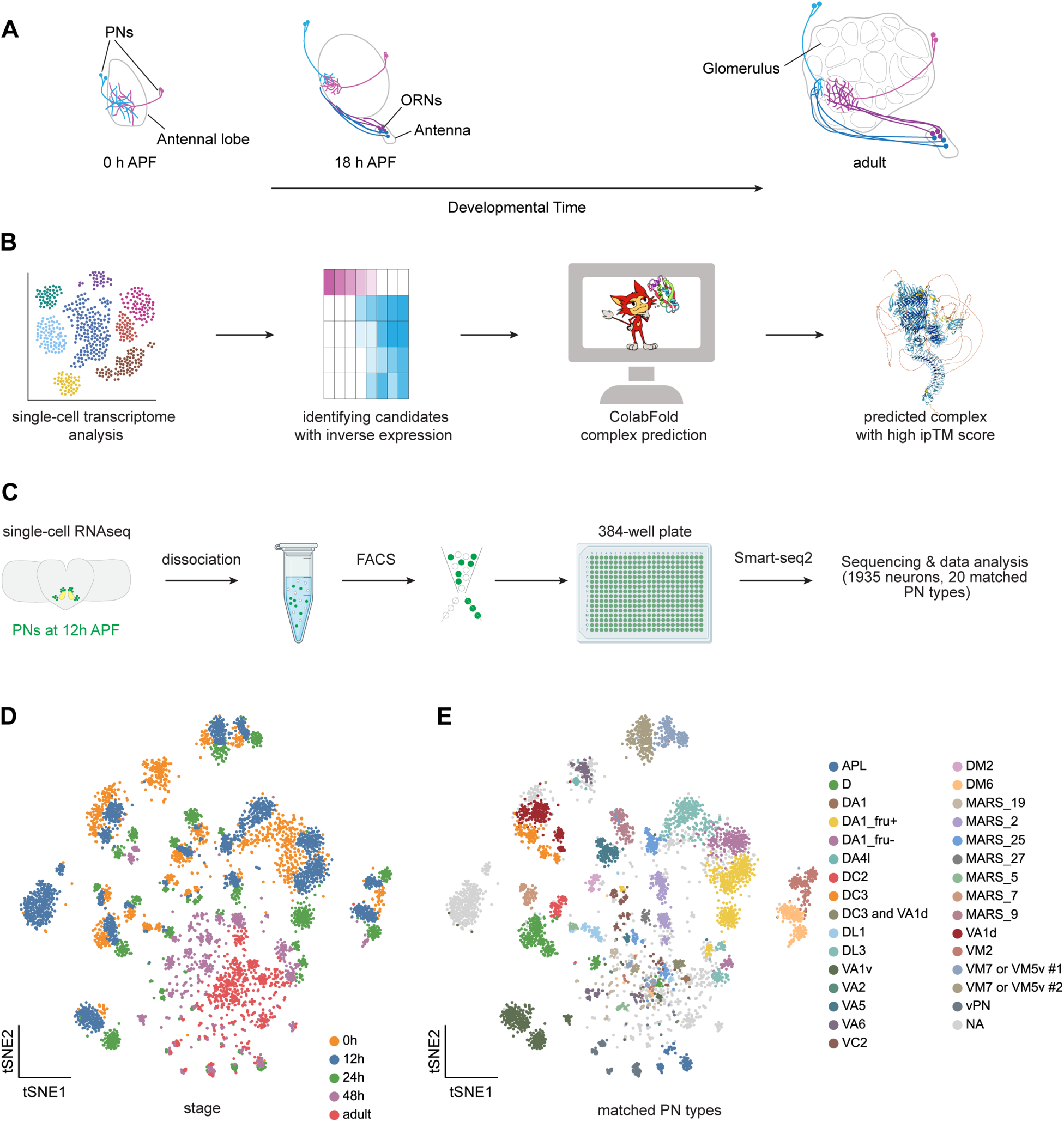
Single-cell transcriptomic profiling of developing *Drosophila* olfactory PNs at 12 h after puparium formation (APF). (A) Schematic of the developing *Drosophila* antennal lobe. PNs, projection neurons. ORNs, olfactory receptor neurons. (B) Schematic of the computational pipeline for identifying Caps binding partners. (C) Workflow for the single-cell RNA sequencing using plate-based SMART-seq2. FACS, fluorescence- activated cell sorting. Dataset integrated with previously published developing PN single-cell transcriptomics^20^ for downstream analyses. (D) Visualization of PN populations across five different developmental stages: 0 h APF, 12 h APF, 24 h APF, 48 h APF, and adult. A total of 545 Iterative Clustering for Identifying Markers (ICIM)^43^ genes at 24 h APF PNs were used for dimensionality reduction. (E) Visualization of the same types of PNs at all developmental stages. Matching colors indicate identical neuronal types; light gray dots, unclassified cells. Cell types lacking definitive identification via cell-type- specific drivers are designated as “MARS_xx” to maintain consistency with previous nomenclature^20^, utilizing the MARS (*meta*-learned representations for single-cell data) annotation method^44^. See Figure S1 for additional data.

The leucine-rich repeat (LRR) transmembrane protein Capricious (Caps) is differentially expressed across PN types and instructs the segregation of Caps-positive and Caps-negative dendrites into discrete, interdigitated glomeruli^9^. Genetic analyses suggest that Caps acts cell-autonomously as a receptor to mediate dendritic segregation by interacting with a heterophilic ligand(s)^9^, but the nature of the ligand(s) is unknown.

Teneurins are an evolutionarily conserved family of type II transmembrane proteins with well-established roles in neural circuit development^10,11^. In the *Drosophila* olfactory system, the two fly teneurins, Ten-m and Ten-a, instruct synaptic partner matching between select ORN–PN pairs through homophilic attraction^12–14^. In the mouse, besides homophilic attraction^15^, reciprocal repulsions between Teneurin-3 (Ten3) and Latrophilin-2 (Lphn2), a member of the adhesion family of G-protein-coupled receptors, also play essential roles in target selection of axons^16–18^. Whether teneurins also participate in heterophilic interactions in neural development in invertebrates has not been reported. Notably, Ten-m has been shown to interact with Tartan (Trn), an LRR protein closely related to Caps, in the regulation of epithelial tissue boundaries^19^, raising the possibility that Ten-m may also interact with Caps.

Here, we leverage single-cell transcriptomic atlases of developing PNs^20^ and AlphaFold2- based structural prediction^21,22^ to identify Ten-m as a heterophilic binding partner of Caps, and validate this binding experimentally. We show that Ten-m and Caps are expressed in largely inverse patterns across PN types during the critical period of dendrite segregation. Through loss-of-function, rescue, and gain-of-function experiments using binding-interface mutants, we demonstrate that heterophilic Ten-m–Caps interactions are required for the segregation of dendrites of different PN types into distinct glomerular territories.

## Results

### Identification of Ten-m as a Caps binding partner

Based on the hypothesis that contact-dependent repulsion drives dendrite segregation during this period, we reasoned that a repulsive ligand for Caps would display an expression pattern inversely correlated with *caps* across PN types. To identify such ligand(s), we developed an integrated pipeline combining single-cell transcriptomics with AlphaFold2-based structural prediction (Figure 1B). We first generated a single-cell transcriptomic atlas of *Drosophila* olfactory PNs at 12–16 hours after puparium formation (h APF), a critical window for dendrite map formation^8^ (Figure 1C). Integration with previously published PN transcriptomes spanning five developmental stages from 0 h APF to adult^20^ yielded transcriptional profiles for diverse PN types across development (Figures 1D and 1E). We then screened for cell-surface proteins (CSPs) with an expression pattern that is anti-correlated with that of *caps* (Figure 2A and Figure S1).

**Figure 2.**
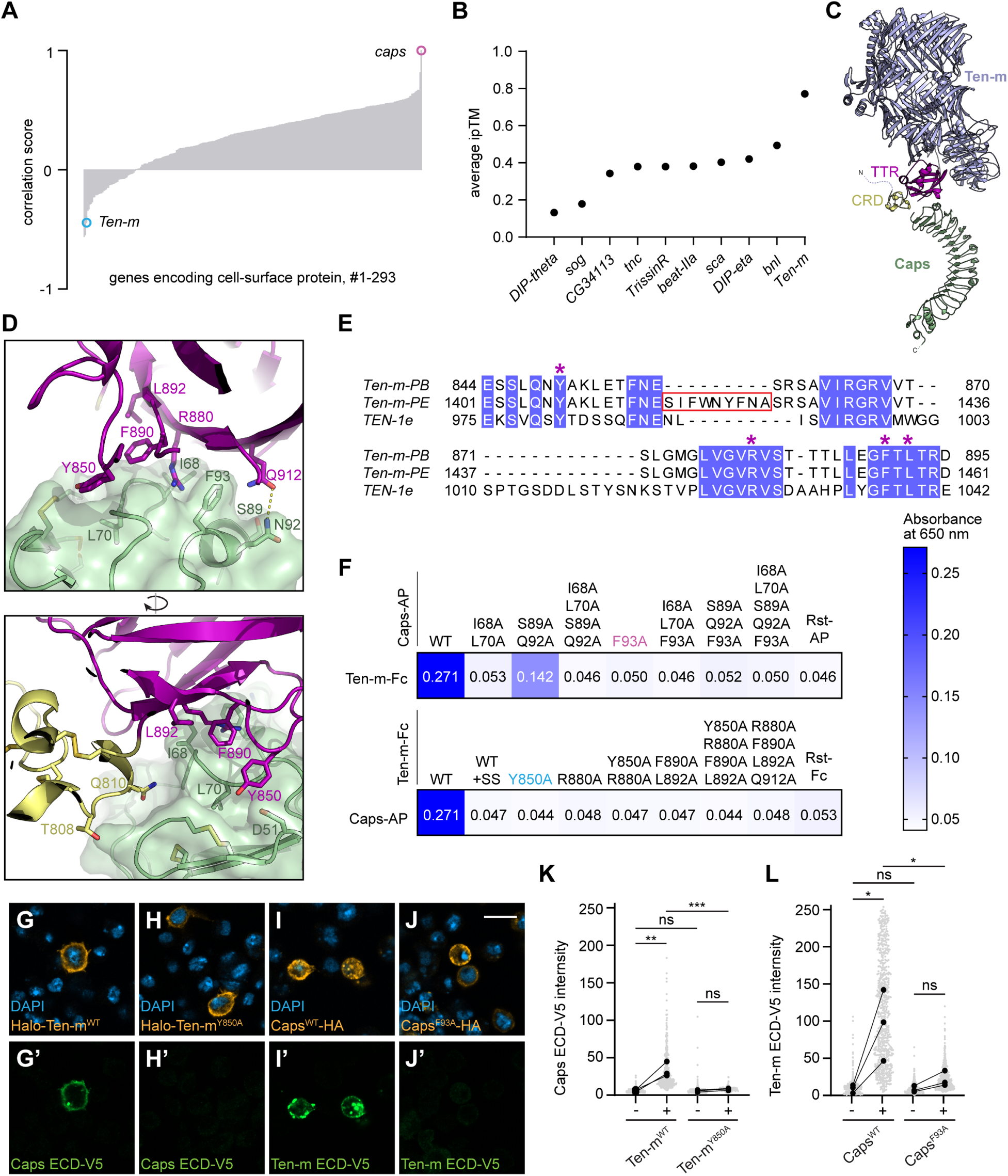
AlphaFold2-guided prediction and experimental validation of the Ten-m–Caps complex binding interface. (A) Correlation between mRNA expression levels of CSPs and *caps* across projection neuron (PN) types, based on single-cell RNA-sequencing data (Figure 1). (B) Ranked ipTM scores for predicted structures formed between the Caps ectodomain (ECD) and ECDs of the top 10 candidate proteins with anti-correlated expression. (C) Structural model of Ten-m (residues 777–2709, isoform B) bound to the Caps ectodomain (green). TTR and CRD domains of Ten-m are colored magenta and yellow, respectively. (D) A detailed view of the residues at the Ten-m–Caps interface with two different viewing angles. Color coding is as in (C). (E) Sequence alignment of TTR domains of Ten-m isoforms B and E, with a splice insertion in E, against *C. elegans* TEN-1. Red box highlights the splice site (+SS) insertion in Ten-m-PE. Stars mark residues chosen for mutation to selectively disrupt the Ten-m–Caps binding. (F) Extracellular interactome assay (ECIA) for Ten-m and Caps mutants designed based on the structural model of the Ten-m–Caps complexes. Fc fusions are immobilized bait, while AP fusions are prey. Colored mutants are used for further *in vitro* and *in vivo* experiments. (G–J’) Caps ECD binds S2 cells expressing Halo-Ten-m^WT^ (G–G’) but not Halo-Ten-m^Y850A^ (H–H’). Ten- m ECD binds S2 cells expressing Caps^WT^-HA (I–I’) but not Caps^F93A^-HA (J–J’). ECDs were visualized with anti-V5 antibodies. Scale bars, 10 μm. (K–L) Quantification of Caps ECD-V5 (K) and Ten-m ECD-V5 (L) binding intensity on cells expressing indicated receptors versus untransfected controls. Repeated measures two-way ANOVA followed by Šídák’s multiple comparisons test (***p < 0.001; **p < 0.01; *p < 0.05; ns, not significant). Gray dots, individual cell intensities; black dots, experimental means (n = 3, 39–985 cells per experiment). Lines connect paired untransfected and transfected populations from the same slide. See Figure S1–S2 for additional data.

From the top 10 anti-correlated candidates, we employed ColabFold (an implementation of AlphaFold2)^23^ to predict interactions between their extracellular domains (ECDs) and the Caps ECD. Ten-m emerged as the leading candidate, forming a predicted complex with Caps with a high inter-chain predicted template modeling (ipTM) score, a metric quantifying confidence in the predicted protein-protein binding interface (Figure 2B).

We find supporting evidence for this interaction from parallel structural studies. In an accompanying paper, we report the cryo-electron microscopy (EM) structure of *C. elegans* TEN- 1 with an LRR domain protein that shares the strongest amino acid sequence similarity with *Drosophila* Caps, LRON-11 (Li et al., companion manuscript), following a high-throughput protein-protein interaction screen^24^. Using this structure as a template for homology modeling, we generated a Ten-m–Caps structural model (Figure 2C) that closely resembled the AlphaFold2 prediction. In both models, the cysteine-rich domain (CRD) and the transthyretin-related (TTR) domain of Ten-m contacted the Caps LRR domain (Figure S2A), supporting our computational approach and suggesting an evolutionary conservation between nematode LRON-11 and *Drosophila* Caps.

Analyzing the modeled Ten-m–Caps interface highlighted a set of conserved residues (Figures 2D and 2E). Using our extracellular interactome assay (ECIA)^25^, we found that Ten-m bound to Caps, and the binding was weakened or abolished by mutations designed at the predicted interface (Figure 2F). Interestingly, an insertion found in one of the Ten-m splice isoforms resulted in loss of binding, suggesting that alternative splicing may play a role in regulating Ten-m interactions with Caps (Figures 2E and 2F). The proper expression and secretion of these mutant ECDs confirmed that loss of binding was specific to interface disruption rather than protein misfolding or trafficking defects (Figures S2B and S2C).

To validate these findings in a cellular context, we performed surface binding assays using *Drosophila* S2 cells^19^. We generated epitope-tagged ECD constructs for both proteins and introduced point mutants designed to disrupt the binding interface. Consistent with ECIA data, S2 cells expressing wild-type Ten-m recruited soluble Caps ECD, whereas S2 cells expressing the binding-deficient Ten-m (Ten-m^Y850A^) failed to do so. Likewise, S2 cells expressing wild-type Caps recruited soluble Ten-m ECD, whereas cells expressing the binding-deficient Caps (Caps^F93A^) failed to do so (Figures 2G–2L). Control experiments confirmed that these mutations did not affect surface trafficking, membrane localization, or neuronal terminal transport of either protein (Figures S2D–S2K’’). Under the same *in vitro* conditions, neither Ten-m nor Caps ECD exhibited robust homophilic binding (Figures S2L–S2O), consistent with a previous report on Ten-m^19^ and suggesting that heterophilic Ten-m–Caps binding is biochemically dominant in isolation. We note that Ten-m homophilic adhesion *in vivo* may require specific membrane microenvironments, clustering, or other factors not captured in our cell-based assay.

Collectively, these data identify Ten-m as a direct binding partner for Caps and support the binding mode predicted by AlphaFold2. Our results demonstrate that the combination of expression analysis and AlphaFold2-based structural prediction can effectively identify binding partners for orphan cell-surface receptors.

### Inverse expression of Ten-m and Caps in the developing antennal lobe

Next, we examined the inverse expression between Ten-m and Caps more closely. At both 12 h and 24 h APF, PN types with high *caps* mRNA expression generally showed low *Ten-m* mRNA expression, and vice versa (Figures 3A and 3B). To validate these findings at the protein level, we took advantage of an endogenously V5-tagged Ten-m line^13^ and generated a Caps knock-in line bearing a C-terminal HA tag. Double immunostaining at 18 h APF, a stage when ORN axons have not yet entered the antennal lobe, revealed a largely inverse expression pattern. Regions enriched for Ten-m were typically devoid of Caps, and vice versa (Figure 3C). Although Ten-m high and Caps-high regions did not strictly follow an anatomical axis, a general trend was that dorsomedial antennal lobe tended to have higher Ten-m expression and ventrolateral antennal lobe had higher levels of Caps. Quantification of fluorescence intensity along this axis validated this trend (Figure 3D).

**Figure 3.**
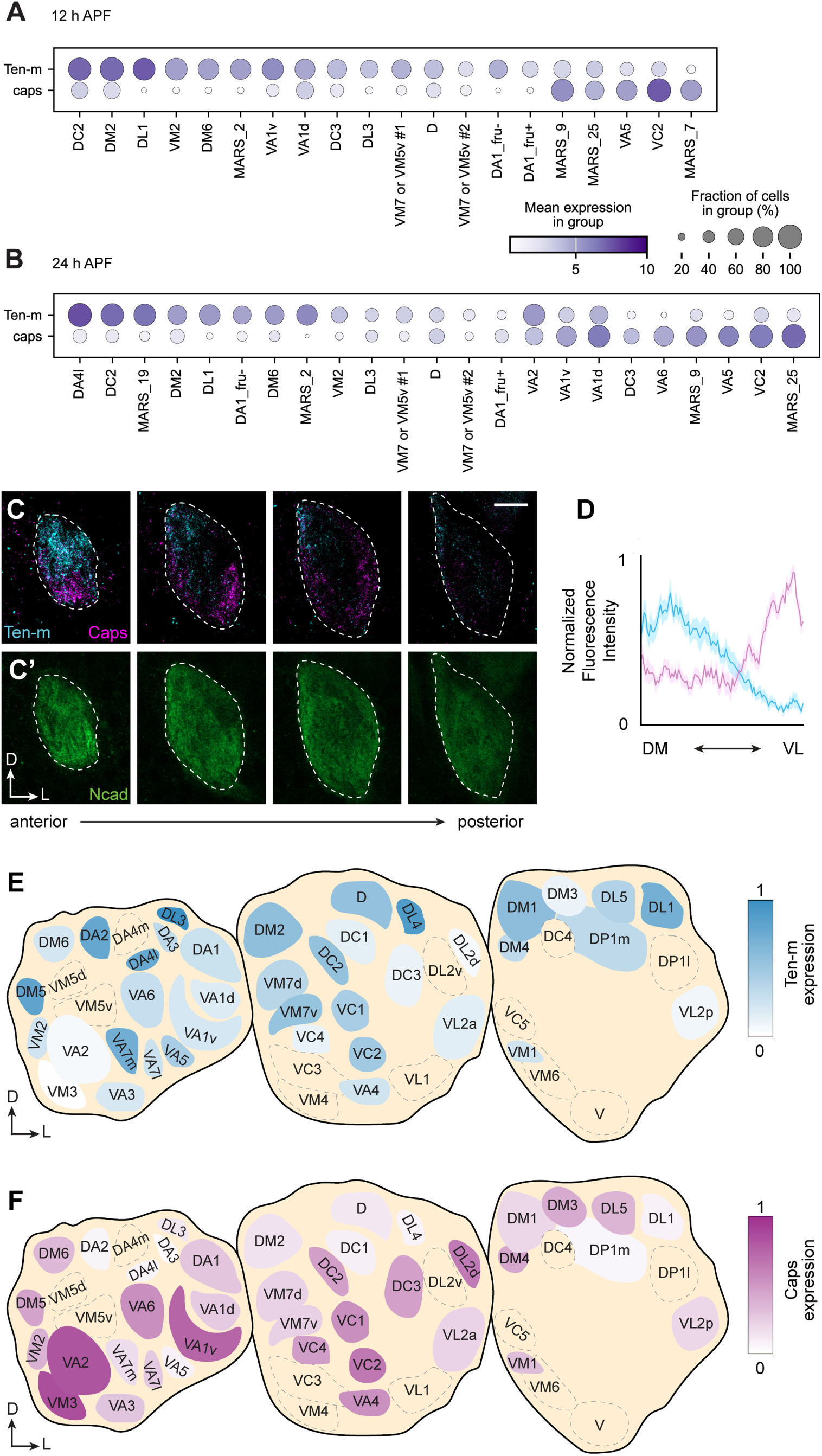
Largely inverse expression patterns of Ten-m and Caps during antennal lobe development. (A, B) Expression profiles of *caps* and *Ten-m* mRNA in PNs at 12 h APF (A) and 24 h APF (B). The unit for “mean expression in group” is log_2_(CPM+1), where CPM is counts per million reads. (C, C’) Double immunostaining of endogenously tagged V5-Ten-m and Caps-HA proteins in the antennal lobe at 18 h APF. Dashed lines indicate antennal lobe boundaries. Scale bars, 10 μm. (D) Quantification of Ten-m and Caps protein distribution along the dorsomedial–ventrolateral (DM–VL) axis of anterior antennal lobe (n = 12 antennal lobes) at 18 h APF. Means ± SEM are shown. (E, F) Normalized protein levels of Ten-m (E; n = 5 antennal lobes) and Caps (F; n = 5 antennal lobes) in PNs across glomeruli at 48 h APF. Dashed areas indicate *GH146*-negative regions where expression was not determined. Endogenous Ten-m and Caps proteins in PNs detected using split-GFP and conditional tagging approaches, respectively. See Figure S3 for additional data. Axis labels in this and all subsequent figures, A, anterior; P, posterior; D, dorsal; V, ventral, M, medial, L, lateral.

Expression pattern analysis after ORN arrival is complicated by the fact that different types of ORNs also differentially express Caps and Ten-m^9,12,26^. To achieve PN-specific protein detection at this stage, we employed a split-GFP-based tag^27^ intersected with a PN-specific driver for Ten- m visualization, and a conditional tag^28^ intersected with PN-specific flippase for Caps detection (Figures S3A and S3B). At 48 h APF, when individual glomeruli become morphologically identifiable, the mutually exclusive expression pattern observed at 18h APF became more intermingled. Although some glomeruli began to co-express intermediate to high levels of both Ten-m and Caps, a substantial portion of the antennal lobe still predominantly conformed to an inverse expression pattern. For instance, the DA2 glomerulus exhibited high Ten-m expression but lacked detectable Caps, whereas the VA2 glomerulus displayed the opposite profile (Figures 3E and 3F). The preferential inverse expression of Ten-m and Caps supports our hypothesis that these proteins define distinct dendritic territories.

Although some glomeruli, such as DA2 and VA2, exhibited clearly distinct Ten-m and Caps levels at 48 h APF, others expressed intermediate levels of one or both proteins. This may be because (1) Ten-m and Caps represent a small fraction of multiple molecular systems that collectively specify ∼50 glomerular territories, and (2) protein expression at this stage may also reflect functions other than dendrite segregation, such as synaptic partner matching. Nevertheless, the glomerulus-level expression map (Figures 3E and 3F) provides a quantitative reference for evaluating the directionality of mistargeting phenotypes in the genetic analyses that follow.

### Both Ten-m and Caps regulate dendrite segregation *in vivo*

To determine if Ten-m and Caps are necessary for dendrite segregation, we employed MARCM (mosaic analysis with a repressible cell marker)^29^ to generate single-cell clones of specific PN types. Because *Ten-m* maps just 0.1 centi-Morgan from the centromere on the chromosomal arm 3L just distal to the insertion site of *FRT2A* commonly used for mosaic analysis^30^, existing alleles cannot be easily recombined onto *FRT2A*. We therefore used CRISPR (clustered regularly interspaced short palindromic repeats) to generate a new *Ten-m* null allele (*Ten-m^Δ^*) directly on a chromosome containing *FRT2A*, deleting exons 3–6 to remove the sole transmembrane domain and introduce a frameshift (Figure S4A).

We focused on DA2-PNs, which exhibit high Ten-m and low Caps expression and target one of the glomeruli with the highest Ten-m levels. In wild-type controls, DA2-PN dendrites were confined to the DA2 glomerulus. We accessed DA2-PNs based on their lineage and birth timing^8^ and confirmed the identity of single-cell clones based on their stereotyped axonal projection patterns^31–33^ (Figure S4B). For each genotype, we scored all identifiable glomeruli innervated by ectopic dendrites and reported those destinations reached in >20% of mutant samples. We found that DA2-PNs homozygous for *Ten-m^Δ^* exhibited severe mistargeting phenotypes, extending dendritic branches into ventral, Caps-high territories (Figures 4A–D), indicating that Ten-m is cell-autonomously required in DA2-PN dendrites to prevent their invasion into Caps-positive territories. Beyond DA2-PNs, several other PN types displayed less pronounced but notable mistargeting to nearby Caps-high glomeruli (Figure S4C), demonstrating that Ten-m function is not restricted to DA2-PNs alone. The specific glomeruli invaded likely reflect the combined influence of earlier patterning cues, such as the semaphorin gradients that constrain dendrites to particular spatial domains^34,35^, and the local availability of Caps-expressing territories along the existing dendritic trajectory.

**Figure 4.**
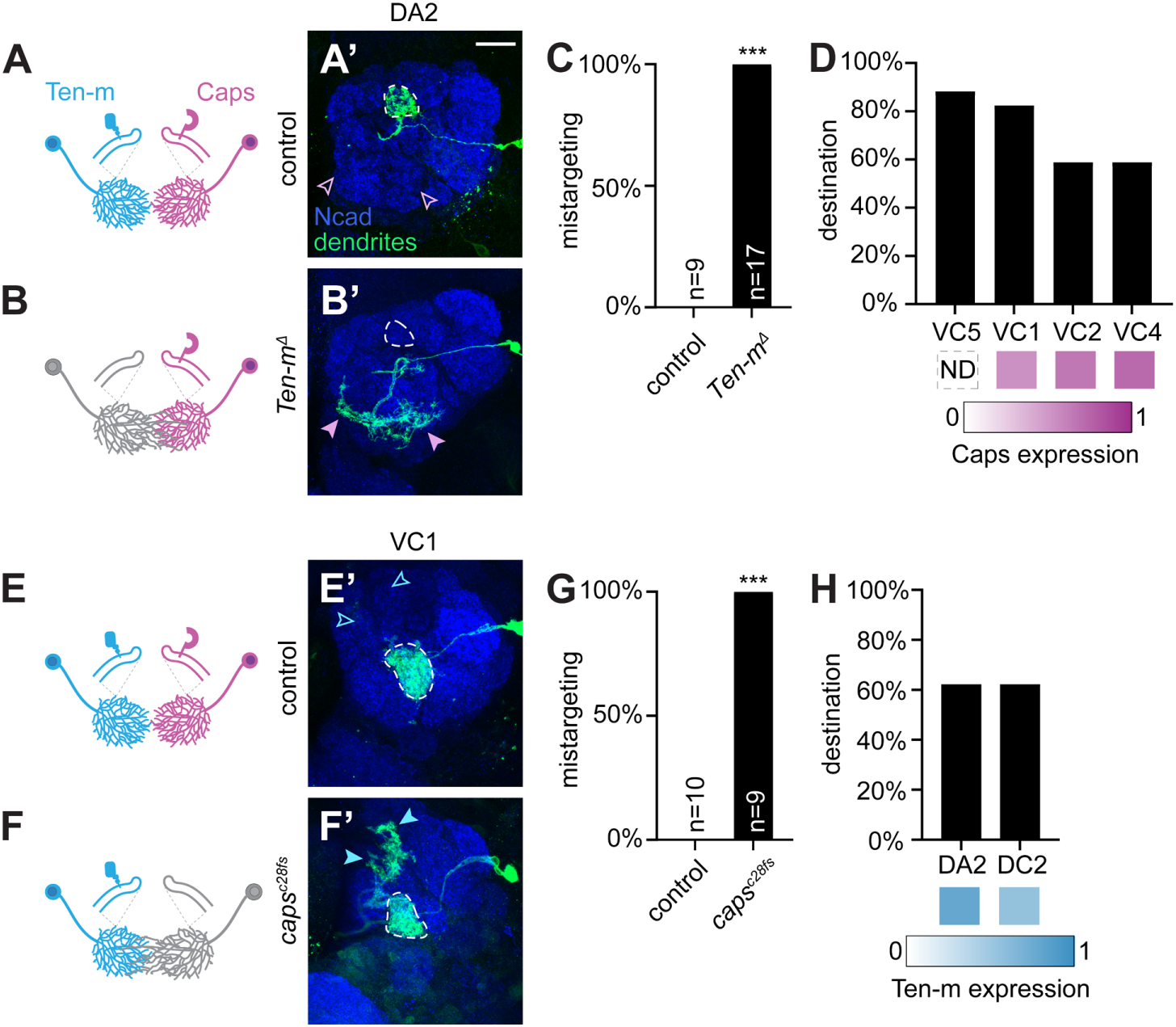
Cell-autonomous requirement of Ten-m and Caps in preventing dendrite invasion into reciprocal expression domains. (A–B’) Schematic and confocal images of dendrites of single DA2-PN MARCM clones for wild-type control (A, A’) or homozygous *Ten-m^Δ^* null allele (B, B’). Dashed lines outline the DA2 glomerulus; open pink arrowheads indicate Caps-high regions normally avoided by DA2-PN dendrites; filled pink arrowheads mark ectopic innervation by *Ten-m^Δ^* dendrites. (C) Phenotypic penetrance of dendrite mistargeting in control and *Ten-m^Δ^*DA2-PN clones. In this and subsequent figures, n represents the number of antennal lobes examined. ***, *p* < 0.001 (Fisher’s Exact Test). (D) Primary mistargeting destinations (>20% penetrance) of DA2-PN dendrites with corresponding Caps expression levels. ND, not determined. (E–F’) Schematic and confocal images of single VC1-PN MARCM clones for control (E, E’) or homozygous *caps^c28fs^* null allele (F, F’). Dashed lines outline VC1 glomerulus; open blue arrowheads indicate Ten-m-high regions normally avoided by VC1-PN dendrites; filled blue arrowheads mark ectopic innervation by mutant dendrites. (G) Phenotypic penetrance of VC1-PN dendrite mistargeting. ***, *p* < 0.001 (Fisher’s Exact Test). (H) Primary mistargeting destinations (>20% penetrance) with corresponding Ten-m expression levels. Scale bar, 20 μm (applies to all confocal image panels). See Figure S4 for additional data.

To determine the developmental timing of Ten-m requirement and distinguish between its roles in dendrite–dendrite segregation versus PN dendrite–ORN axon matching, we examined DC2-PNs, for which we have a specific driver, allowing us to examine dendritic development of a specific PN type at 18 h APF, prior to the onset of dendrite–axon matching. Whereas control DC2-PN dendrites remained confined to a dorsal region, Ten-m knockdown resulted in dendrite bifurcation and aberrant extension into the ventral, Caps-high region of the antennal lobe (Figures S4D and S4E). These findings suggest that Ten-m is required for proper DC2-PN dendrite targeting from early developmental stages.

We next examined the cell-autonomous function of Caps in dendrite segregation focusing on VC1-PNs, which normally express high Caps and low Ten-m levels. Consistent with previous reports^9^, loss of *caps* in VC1-PNs caused dendrite invasion into ectopic regions. Importantly, these ectopic targets were predominantly Ten-m-high glomeruli (Figures 4E–4H).

To test whether *Ten-m* and *caps* genetically interact during dendrite segregation, we examined DL3-PN dendrite targeting in *Ten-m*/*caps* trans-heterozygotes (losing one copy of both genes). DL3-PNs innervate a glomerulus with high Ten-m and low Caps expression, adjacent to the DA1 glomerulus, which exhibits low Ten-m and moderate Caps levels. We took advantage of a driver that selectively labels DL3-PNs at the adult stage to achieve sparse labeling of individual DL3-PNs. Losing one copy of *Ten-m* or *caps* alone did not cause dendrite targeting deficits; however, trans-heterozygotes exhibited significantly increased dendrite mistargeting (Figures S4F and S4G). These data are consistent with *Ten-m* and *caps* working in the same pathway.

A key observation from both *Ten-m* and *caps* loss-of-function experiments is that dendrite mistargeting was not random but directed toward specific glomeruli. This specificity argues against a simple loss of homophilic adhesion among dendrites of the same PN type. Instead, Ten-m-expressing dendrites invaded Caps territories upon Ten-m loss and Caps-expressing dendrites invaded Ten-m territories upon Caps loss. These phenotypes support a model in which heterophilic Ten-m–Caps interactions mediate mutual repulsion between dendrites of Caps-high and Ten-m- high PNs, thereby facilitating their segregation.

### Ten-m–Caps binding is required for dendrite segregation

We next tested whether the physical interaction between Ten-m and Caps is necessary for their function in dendrite segregation using the binding-deficient mutants (Figure 2). Teneurins are well-established mediators of synaptic partner matching through homophilic attraction^12,13,15,36^. Structural analysis of the Ten-m dimer reveals that the CRD-TTR interface, which is critical for Ten-m–Caps interaction, also mediates Ten-m–Ten-m homophilic interaction^37^. We therefore asked whether the Caps-binding-deficient *Ten-m^Y850A^* mutation affects Ten-m’s homophilic function in synaptic partner matching.

To test this, we overexpressed *Ten-m^WT^* and *Ten-m^Y850A^*in DA1-ORNs. Consistent with previous work showing that *Ten-m* overexpression in ORNs induces mismatching with PNs through homophilic attraction^13^, we found that *Ten-m^Y850A^* induced ORN–PN mismatching as effectively as the wild-type *Ten-m* (Figures S5A–S5D). This result suggests that the Y850A mutation preserves Ten-m homophilic interaction while disrupting Caps binding, allowing us to assess specifically the function of Ten-m–Caps interaction using this allele.

We thus introduced the *Ten-m^Y850A^* mutation into the endogenous locus through CRISPR knockin^38^. PNs homozygous mutant for *Ten-m^Y850A^* recapitulated the dendrite segregation defects observed in *Ten-m^Δ^* null mutants (Figures 5A–5D). Altogether, these findings suggest that heterophilic binding to Caps supports Ten-m’s role in dendrite segregation, independent of Ten- m’s roles in synaptic partner matching supported by homophilic Ten-m interaction.

**Figure 5.**
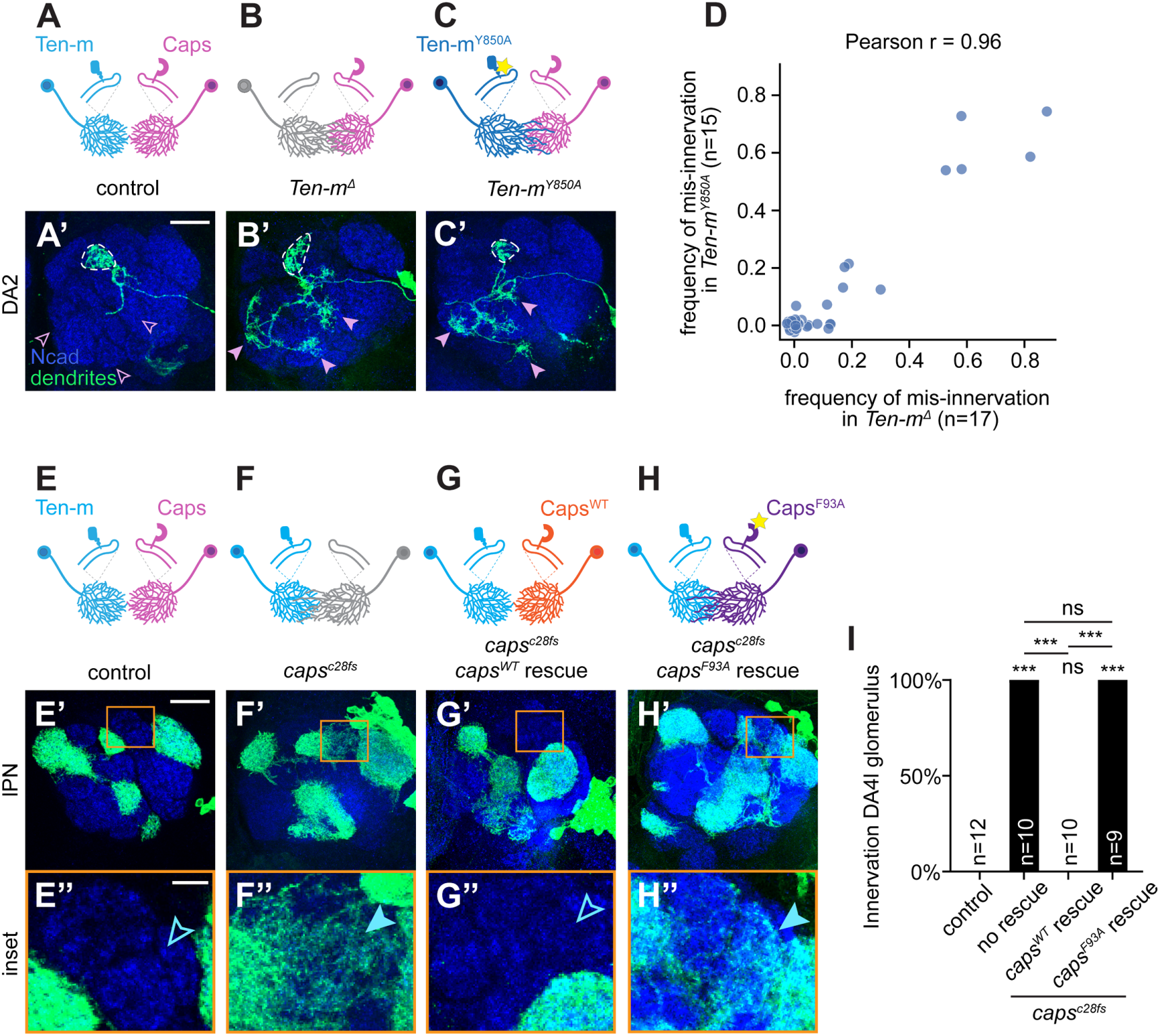
Ten-m–Caps binding is essential for dendrite segregation. (A–C’) Schematics and confocal images of dendrites of single DA2-PN MARCM clone with wild-type (A, A’), *Ten-m^Δ^* null allele (B, B’), or *Ten-m^Y850A^* encoding Caps-binding-deficient Ten-m (C, C’). Dashed lines outline the DA2 glomerulus; open/filled pink arrowheads indicate Caps-high regions normally avoided or ectopically innervated, respectively. (D) Comparison of mis-innervation frequencies between *Ten-m^Y850A^* and *Ten-m^Δ^* DA2-PN dendrites. Each dot represents a non-DA2 glomerulus; the x- and y-axis values represent the frequencies of dendrite mis- innervation by DA2-PNs mutant for *Ten-m^Δ^*and *Ten-m^Y850A^*, respectively. (E–H’’) Schematics and confocal images of lateral PN (lPN) neuroblast MARCM clones with wild-type *caps* (E–E’’), *caps^c28fs^* (F–F’’), *caps^c28fs^*rescued with *caps^WT^* (G–G’’), or *caps^c28fs^* rescued with *caps^F93A^* encoding Ten-m–binding-deficient Caps (H–H’’). Open/filled blue arrowheads indicate Ten-m-high regions normally avoided or ectopically innervated, respectively. (I) Quantification of ectopic DA4l innervation across genotypes. Statistical differences between conditions were assessed using 2×2 Fisher’s Exact Test followed by Benjamini-Hochberg (FDR) correction for all pairwise comparisons. ns (not significant); ***, adjusted p < 0.001. Statistical comparisons shown directly above each bar are relative to the control group. Scale bar shown in (A’ and E’): 20 μm, applicable to panels (B’, C’, F’, G’, and H’). Scale bar shown in (E’’): 5 μm, applicable to panels (F’’, G’’, and H’’). See Figure S5 for additional data.

We similarly examined whether Ten-m binding is required for Caps function in dendrite segregation by employing a transgene rescue strategy. Unlike wild-type lateral neuroblast MARCM clones (Figure 5E–5E’’), in lateral neuroblast clones homozygous mutant for a *caps* null mutant, PN dendrites mistargeted to regions normally occupied by PNs from the anterodorsal lineage, including the DA4l glomerulus (Figure 5F–5F’’). Expression of wild-type *caps* in *caps* null lateral PN MARCM neuroblast clones rescued dendrite targeting phenotypes (Figure 5G–5G’’). However, expressing the Ten-m-binding-deficient *caps^F93A^* mutant failed to rescue (Figure 5H–5H’’, quantified in Figure 5I). These experiments demonstrated that Ten-m binding is essential for Caps function.

Two gain-of-function experiments further supported the repulsive interaction model. In the first experiment, we misexpressed *caps* in Ten-m-high/Caps-low DL1-PNs and found that this caused dendrite mistargeting to Ten-m-low/Caps-high regions; however, similar misexpression of *caps^F93A^*showed significantly reduced mistargeting phenotypes (Figures S5E–S5H), supporting that Caps–Ten-m interaction is required for this gain-of-function phenotype. In the second experiment, we misexpressed *Ten-m* in Mz19-PNs (which innervate DA1, VA1d, and DC3 glomeruli—with DA1-PNs and DC3-PNs normally expressing low Ten-m^12^) and found that this caused their dendrites to shift away from their normal target and invade Ten-m-high/Caps-low regions (Figures S5I–S5L). Notably, misexpression of the non-binding mutant *Ten-m^Y850A^* resulted in dendrite invasion of the Caps-high VA1v glomerulus, a phenotype not observed with wild-type Ten-m misexpression (Figures S5J–S5K). That the specific mistargeting destinations differ between wild-type and binding-deficient variants indicates that Ten-m–Caps binding biases which regions dendrites avoid. The observation that misexpression of *caps^F93A^* or *Ten-m^Y850A^*still exhibited mistargeting phenotypes suggests either that these mutations did not eliminate binding, or that misexpressed *caps^F93A^*or *Ten-m^Y850A^* may engage other interactions to drive mistargeting.

Results from the loss-of-function, rescue, and gain-of-function experiments collectively demonstrate that the Ten-m–Caps binding interface identified by structural modeling is functionally required *in vivo* to enforce dendrite segregation through heterotypic repulsion.

## Discussion

In this study, we show that heterophilic binding between Ten-m and Caps drives the segregation of PN dendrites into discrete glomerular territories in the *Drosophila* antennal lobe. Our findings establish a molecular mechanism for heterotypic dendrite segregation—a developmental process that is critical for the formation of discrete neural maps.

### Reciprocal repulsion for dendrite segregation

Previous work suggested a hierarchical model for PN dendrite targeting in which global semaphorin gradients first define coarse spatial domains, followed by local segregation into discrete glomeruli mediated by molecules such as Caps^9,34,35^. However, how Caps mediates local segregation has been unclear, as it does not function homophilically and its binding partner was unknown. Our identification of Ten-m as the Caps partner fills this gap. The cell-autonomous loss-of-function phenotypes for both Ten-m and Caps support a model in which their binding leads to mutual repulsion between Ten-m^+^ and Caps^+^ dendrites. This property, along with the largely complementary expression of Ten-m and Caps, provides an intuitive solution to the problem of heterophilic dendrite segregation, enforcing mutual exclusion between molecularly distinct dendritic populations. How trans-cellular binding between Ten-m and Caps translates into repulsion at the cellular level remains to be determined. We speculate that contact between Ten- m- and Caps-expressing dendrites triggers intracellular signaling that causes dendrites to withdraw, analogous to other contact-dependent repulsion systems in which receptor binding initiates cytoskeletal remodeling^39^. The mechanism by which Ten-m–Caps binding triggers repulsion remains to be investigated.

This contact-dependent heterophilic repulsion between inversely expressed cell-surface proteins is mechanistically distinct from the self-avoidance pathways mediated by Dscam or protocadherins, which use homophilic interactions to repel sister branches of the same neuron^3,40^. Whereas self-avoidance ensures uniform coverage of a territory by one neuron’s dendrites, the heterophilic repulsion we describe here partitions shared territory between neurons of different types.

This heterophilic exclusion logic may operate more broadly across other parts of the nervous system and in other organisms. Specifically, it could provide a general mechanism for interdigitated neuronal types to establish and maintain discrete territories. For example, in the vertebrate retina, repulsive signaling between differentially expressed transmembrane semaphorins and their plexin receptors restricts the neurites of specific amacrine and retinal ganglion cells to distinct sublaminae^41^. Similarly, in the mouse hippocampal network, Ten3 and Lphn2 are expressed in opposing patterns, where contact-dependent mutual repulsion between them instructs the precise target selection of projecting axons^16–18^. While these studies demonstrate that heterophilic repulsion can pattern axonal targeting and neurite stratification, it has not been shown whether similar mechanisms drive a distinct developmental problem: the segregation of dendrites from different neuronal types into discrete territories. Our results provide evidence that such heterotypic dendrite segregation can be enforced by contact-dependent heterophilic repulsion between inversely expressed cell-surface proteins. Whether this principle extends to vertebrate circuits with similar inversely expressed receptor-ligand pairs remains to be explored.

### Distinguishing heterophilic from homophilic contributions at a shared interface

Because homophilic (Ten-m–Ten-m) and heterophilic (Ten-m–Caps) binding both require the CRD-TTR region^37^, a key question is whether the dendrite segregation phenotypes reflect loss of one interaction, the other, or both. Several lines of evidence support a primary role for heterophilic binding in dendritic segregation. First, the *Ten-m^Y850A^* mutation that abolishes Caps binding retains homophilic function—at least in an overexpression-based synaptic partner matching assay—yet phenocopies the *Ten-m* null for dendrite segregation. Second, on the Caps side, the Ten-m-binding- deficient *caps^F93A^*fails to rescue dendrite targeting in *caps* null clones, independently demonstrating that the Ten-m–Caps interaction is required for Caps function. Third, the trans-heterozygous genetic interaction between *Ten-m* and *caps* provides another evidence that the two genes function in the same pathway. Fourth, the directional specificity of mistargeting—Ten-m- positive dendrites invading Caps-positive territories and vice versa—is most consistent with loss of a heterophilic repulsion rather than loss of homophilic adhesion.

We note, however, that because the two interfaces overlap, we cannot entirely exclude a partial contribution of reduced homophilic binding at endogenous expression levels for the *Ten- m^Y850A^* allele. Future identification of mutations that selectively disrupt Ten-m homophilic but not heterophilic binding or a comprehensive interactome analysis of Ten-m and Caps binding partners will be valuable for fully delineating the network of interactions that shape the glomerular map. Nevertheless, the convergent evidence discussed above makes it unlikely that both coincidentally disrupt unrelated interactions and thus strongly support heterophilic Ten-m–Caps interaction as a major driver of dendrite segregation.

### Co-expression and potential *cis*-regulation

Although Ten-m and Caps show predominantly inverse expression, some PN types co-express both proteins, raising the question of how the two interact when present on the same cell surface—for example via *cis*-inhibition that renders such neurons insensitive to *trans* signaling, with relative levels or subcellular localization tuning the balance. Relatedly, our finding that an alternatively spliced insertion in Ten-m abolishes Caps binding suggests that splicing may regulate the choice between Ten-m’s homophilic (synaptic partner matching) and heterophilic (dendrite segregation) functions in a cell-type-specific manner. Future work mapping the *cis*/*trans* logic and the splice-isoform landscape across PN types will be needed to test these hypotheses.

### Methodological implications

Our identification of the Ten-m–Caps interaction illustrates a generalizable pipeline for de-orphaning ligand–receptor interactions by integrating single-cell transcriptomics, structural prediction, structure-guided mutagenesis, and *in vivo* functional validation. Anti-correlated expression across PN types narrowed the candidate pool, and ColabFold-based complex prediction identified Ten-m as the most structurally plausible interactor. The predicted binding mode was independently validated by the cryo-EM structure of the orthologous *C. elegans* TEN-1/LRON-11 complex (Li et al., companion manuscript), and the predictions enabled design of binding-interface mutations that allowed direct *in vivo* testing—a workflow that would be substantially slower using biochemical approaches alone. Although we used ColabFold/AlphaFold2 for its scalability, AlphaFold3^42^ offers enhanced accuracy by modeling post-translational modifications; our independent validation with AlphaFold3 confirmed the predicted Caps–Ten-m binding mode. Combined with the rapid expansion of single-cell atlases and the increasing accuracy of complex prediction tools, this integrated approach should accelerate the discovery of functionally important cell-surface interactions across biological systems.

## Resource Availability Lead Contact

Further information and requests for resources and reagents should be directed to and will be fulfilled by the lead contact, Liqun Luo.

## Materials Availability

All new DNA constructs generated in this study will be deposited in Addgene, and new transgenic flies generated in this study will be deposited in Bloomington *Drosophila* Stock Center. Other reagents are available from the Lead Contact upon request.

## Data and Code Availability

- Single-cell RNA-seq data generated in this study have been deposited in the NCBI Gene Expression Omnibus (GEO). Accession number: GSE318611. This paper also analyzes existing, publicly available data: GSE161228^20^.
- All original code for single-cell RNA-seq analysis and anti-correlation analysis has been deposited on GitHub (https://github.com/phagocyt/FlyPN).
- Any additional information required to reanalyze the data reported in this paper is available from the lead contact upon request.

## Acknowledgments

We thank the Bloomington *Drosophila* Stock Center and Vienna *Drosophila* Resource Center for fly stocks; J. Zallen for *Ten-m* and *trn*-expressing constructs; K. Shen and members of the Luo laboratory, especially C. McLaughlin, D. Pederick, T. Hindmarsh Sten, J. Kalai and R. Du, for discussions. Computational resources were provided by the Sherlock and Marlowe high-performance computing clusters at Stanford University. This work was supported by the National Institutes of Health grants R01-DC005982 (L.L.) and R35-GM148412 (to D.A.). H.J. was supported by the American Heart Association Postdoctoral Fellowship (24POST1186864/2024). L.L. is an investigator of the Howard Hughes Medical Institute.

## Author Contributions

H.J. and L.L. conceived the project, designed the experiments, interpreted the data, and wrote the manuscript with input from all authors. K.K.L.W. and Y.W. performed the scRNA-seq experiments with support from Z.L., J.L., R.C. J., and S.R.Q. Single-cell transcriptomic data were analyzed by Y.W., K.K.L.W., and H.J. H.J. conducted the *in-silico* AF-M screening. J.L., D.A., and E.Ö. constructed the structural models and performed the biochemical assays. H.J., Y.X., and Y.Z. conducted the *in vivo* experiments, with D.J.L. assisting in the generation of transgenic flies.

---

**Figure S1.**
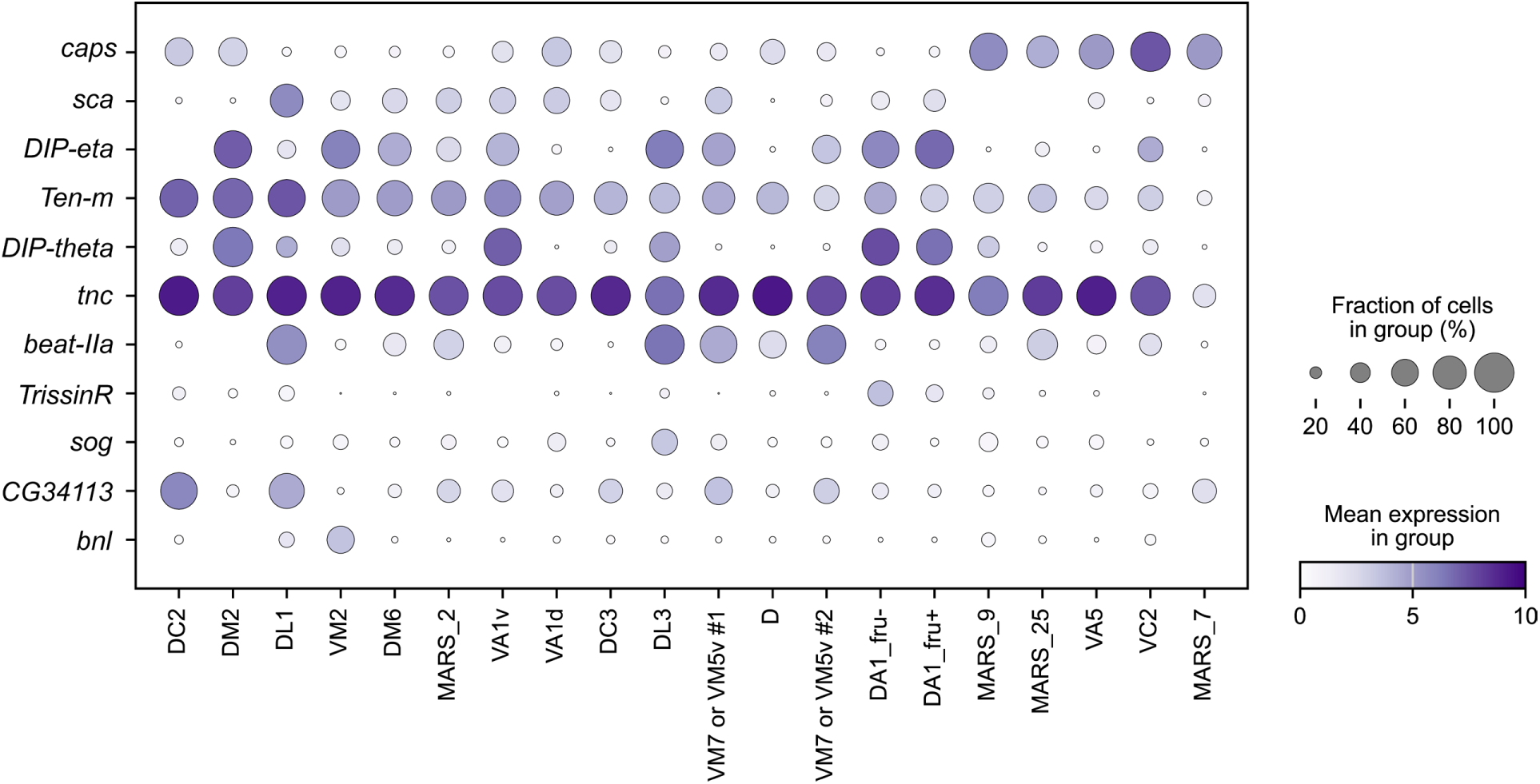
Top 10 candidate cell-surface protein genes with expression anti-correlated to *caps* in projection neurons at 12 h APF, related to Figure 1 and Figure 2. Expression profiles of *caps* and top 10 anti-correlated genes at 12 h APF, shown across PN types. Dot size, percentage of expressing cells; color intensity, mean expression level [log_2_(CPM+1), where CPM is counts per million reads]. The *caps* and *Ten-m* rows are reproduced as Figure 3A.

**Figure S2.**
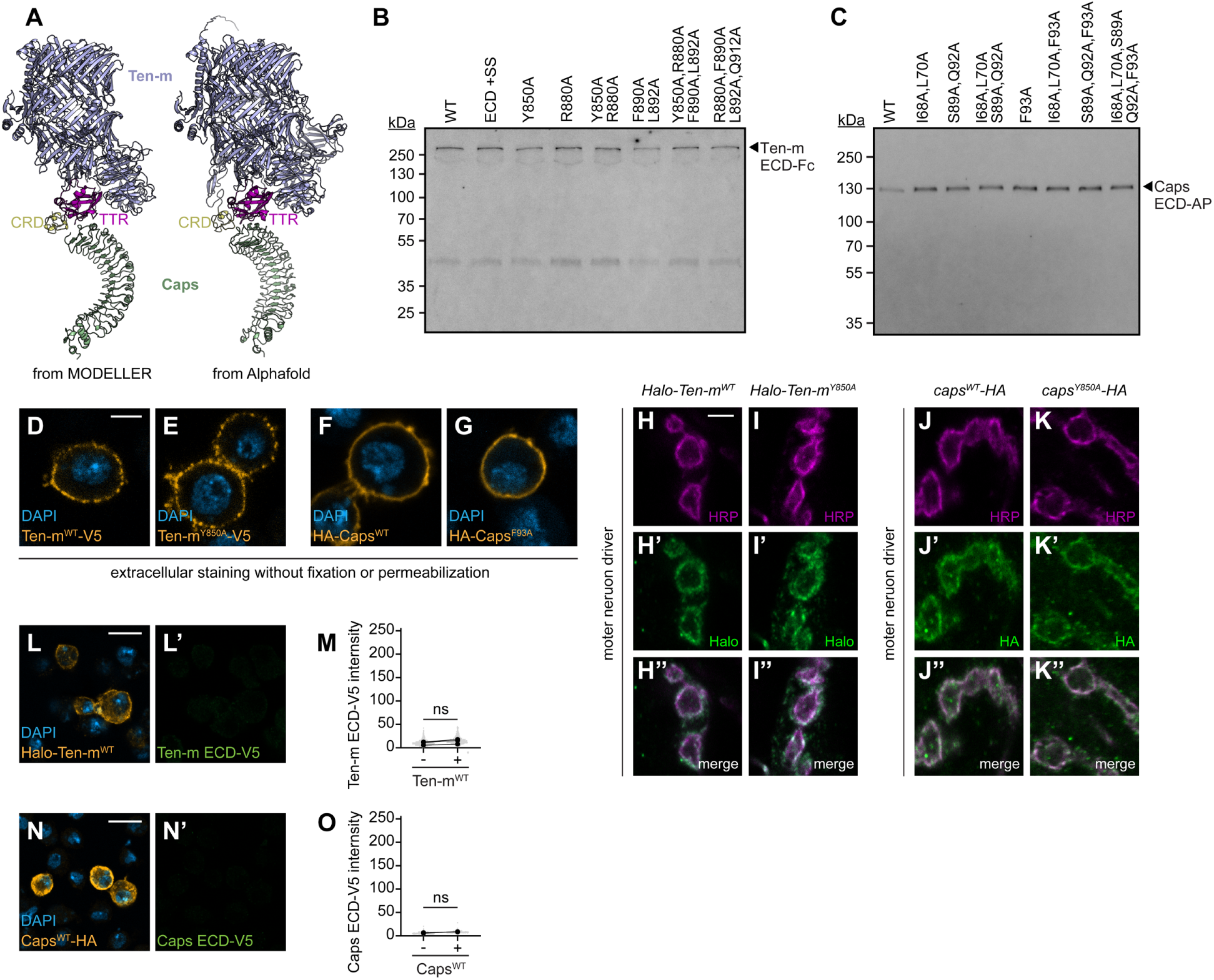
Expression and subcellular localization of Ten-m and Caps variants, related to Figure 2. (A) Model of Ten-m–Caps complex, derived via homolog modeling from our experimentally determined TEN-1–LRON-11 structure (left), closely resembles the AlphaFold2-derived Ten-m–Caps model (right). Low confidence (low pLDDT, predicted local distance difference test) residues were hidden in the AlphaFold2 model. (B, C) Western blots of media of cultured cells expressing ECDs of Ten-m and Caps show that Ten-m and Caps mutant ECDs express and secrete, which are then used in the ECIA. Molecules were detected using an anti-pentaHis antibody that recognizes both Fc- and AP-tagged constructs via a common C-terminal hexahistidine tag. (D–G) Surface expression of Ten-m and Caps variants in S2 cells. Live staining of extracellular epitopes confirms proper membrane trafficking: Ten-m^WT^-V5 (D), Ten-m^Y850A^-V5 (E), HA-Caps^WT^ (F), or HA- Caps^F93A^ (G). Scale bar, 3 μm. (H–K’’) Neuromuscular junction localization of intracellularly tagged constructs expressed via *D42- GAL4* expressed in motor neurons. All variants (labeled on top) colocalize with neuronal membranes at HRP-positive synaptic boutons at the neuromuscular junction, confirming protein stability and axonal transport *in vivo*. Scale bar, 3 μm. (L–L’, N–N’) Cell-based binding assays testing homophilic interactions. S2 cells expressing Halo-Ten- m^WT^ incubated with soluble Ten-m ECD-V5 (L–L’) and S2 cells expressing Caps^WT^-HA incubated with soluble Caps ECD-V5 (N–N’). Scale bars, 10 μm. (M, O) Quantification of Ten-m ECD-V5 (M) and Caps ECD-V5 (O) binding intensity on cells expressing indicated receptors versus untransfected controls. Paired *t*-test (ns, not significant). Gray dots, individual cell intensities; black dots, experimental means (n = 3, 70–909 cells per experiment). Lines connect paired untransfected and transfected populations from the same slide.

**Figure S3.**
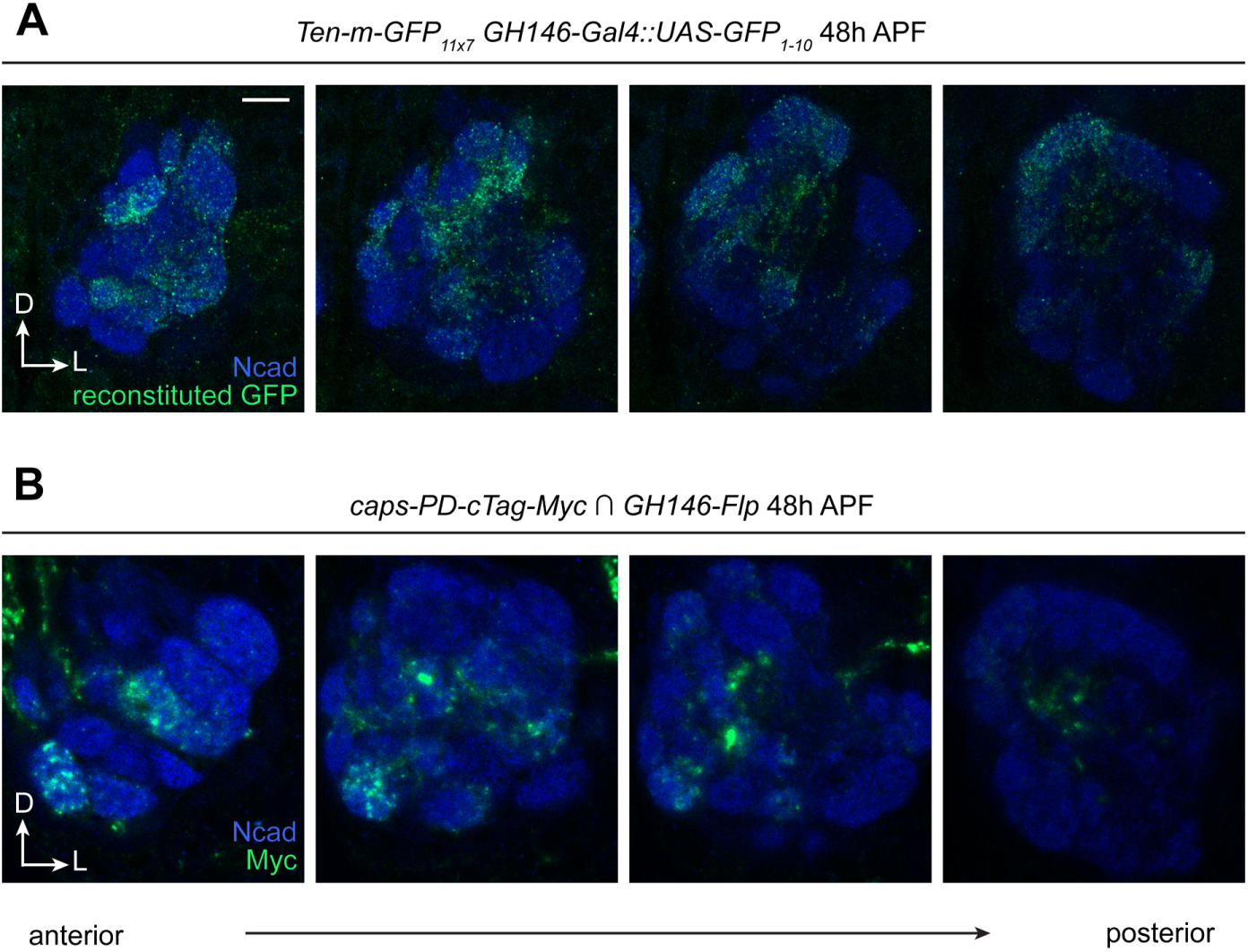
Endogenous Ten-m and Caps expression patterns in PNs at 48 h APF, related to Figure 3. (A) Split-GFP visualization of endogenous Ten-m proteins in PNs. Ten-m is endogenously tagged with *GFP_11×7_*^27^; *GFP_1–10_* is expressed under *GH146-GAL4* driver. Reconstituted GFP detected by immunostaining with a conformation-specific anti-GFP antibody. (B) Conditional tagging strategy for PN-specific Caps visualization using *Caps-PD-HA>stop>Myc* intersected with *GH146-Flp*, detected via anti-Myc immunostaining. Both panels show a representative antennal lobe in four single confocal sections from anterior to posterior. Scale bars, 10 μm. Note that *GH146* drivers are expressed in most but not all PN types, limiting detection coverage.

**Figure S4.**
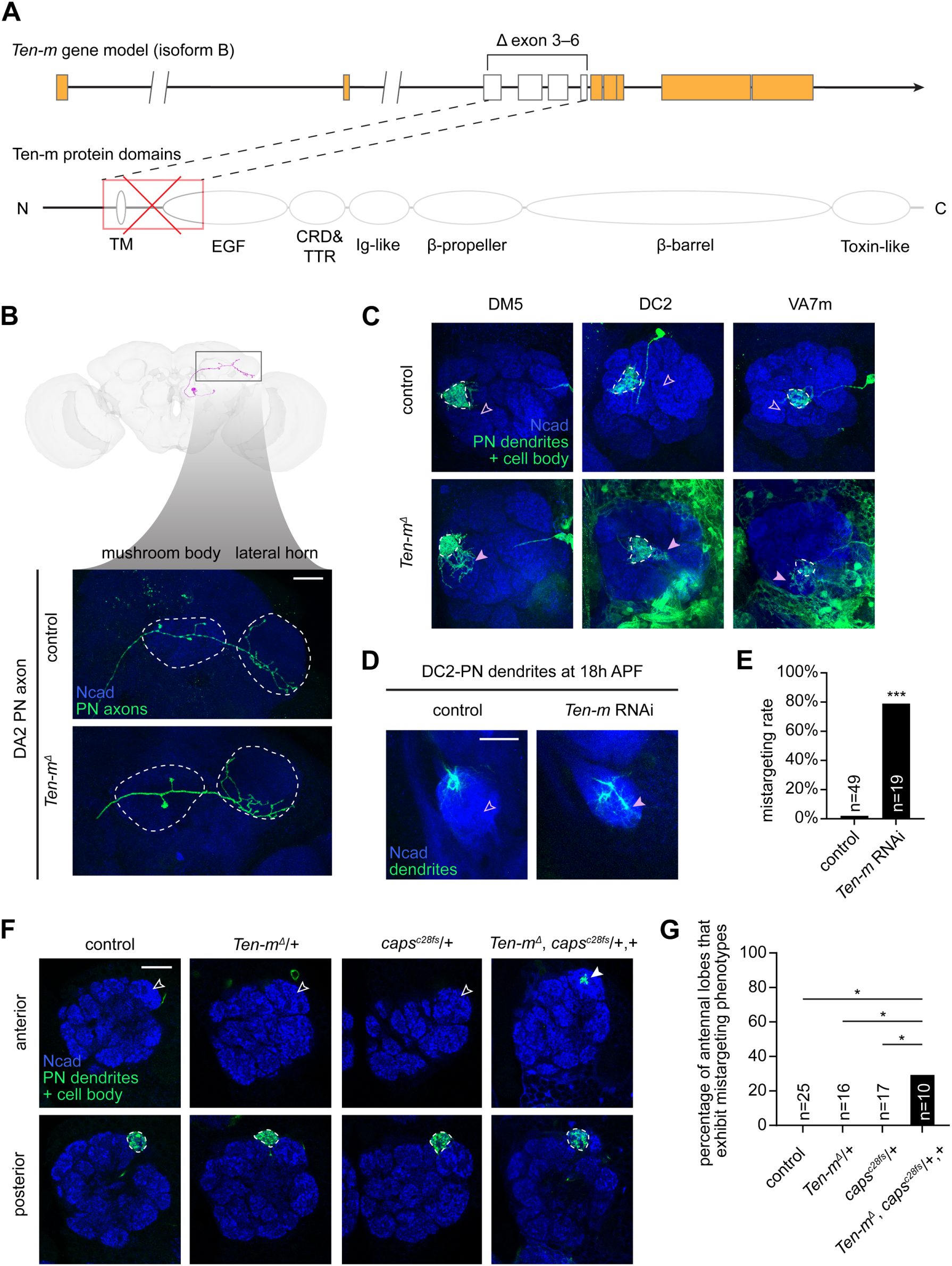
*In vivo* characterization of *Ten-m* loss-of-function and its genetic interaction with *caps*, related to Figure 4. (A) Top: Schematic of the *Ten-m^Δ^*allele showing the gene model of isoform B (filled boxes, coding exons; lines, introns; exons 3–6 shown as open boxes within the bracket-marked deletion). Double slashes (//) indicate a large intron drawn not to scale. Bottom: Ten-m protein domain architecture (isoform B) with major domains depicted as labeled shapes. Dashed lines connect the deleted exons to the TM-encoding region of the protein. The deletion eliminates the TM domain (red cross) and the resulting frameshift truncates all downstream extracellular sequence (shown faded), generating a predicted null protein product. The deletion is predicted to disrupt all Ten-m isoforms. (B) Tracings of a DA2-PN from the FlyWire dataset^31–33^; boxed region indicates axon branching in the mushroom body and lateral horn (top). Representative confocal images of DA2-PN axon morphology in control and *Ten-m^Δ^* mutants. Mutant clones were unambiguously identified as DA2-PNs based on their stereotyped axonal arborization in the lateral horn. This morphology is clearly distinct from that of DM5- PNs, the other principal lateral PN generated during the same 96–120 h heat-shock window. (C) Examples of dendrite mistargeting in *Ten-m^Δ^* PNs. Dashed lines, native glomerular targets; open/filled pink arrowheads, normally uninnervated/ectopically innervated regions. (D) DC2-PN dendrite mistargeting upon *Ten-m* knockdown at 18–21 h APF. Dorsal-to-ventral shift indicated by empty/filled pink arrows. (E) Quantification of DC2-PN mistargeting. 2×2 Fisher’s Exact Test; ***, p < 0.001. (F) DL3-PN dendrite mistargeting in DA1 glomerulus in *Ten-m*/*caps* trans-heterozygotes, but not in wild-type controls or single-heterozygotes. Dashed lines outline DL3 glomerulus; open/filled arrowheads, normally uninnervated/ectopically innervated regions. (G) Quantification of percentage of antennal lobes that exhibit mistargeting phenotypes across genotypes. Statistical differences between conditions were assessed using 2×2 Fisher’s Exact Test followed by Benjamini-Hochberg False Discovery Rate (FDR) correction for 3 pre-planned comparisons (trans- heterozygotes vs. control and each single heterozygote). *, adjusted p < 0.05.

**Figure S5.**
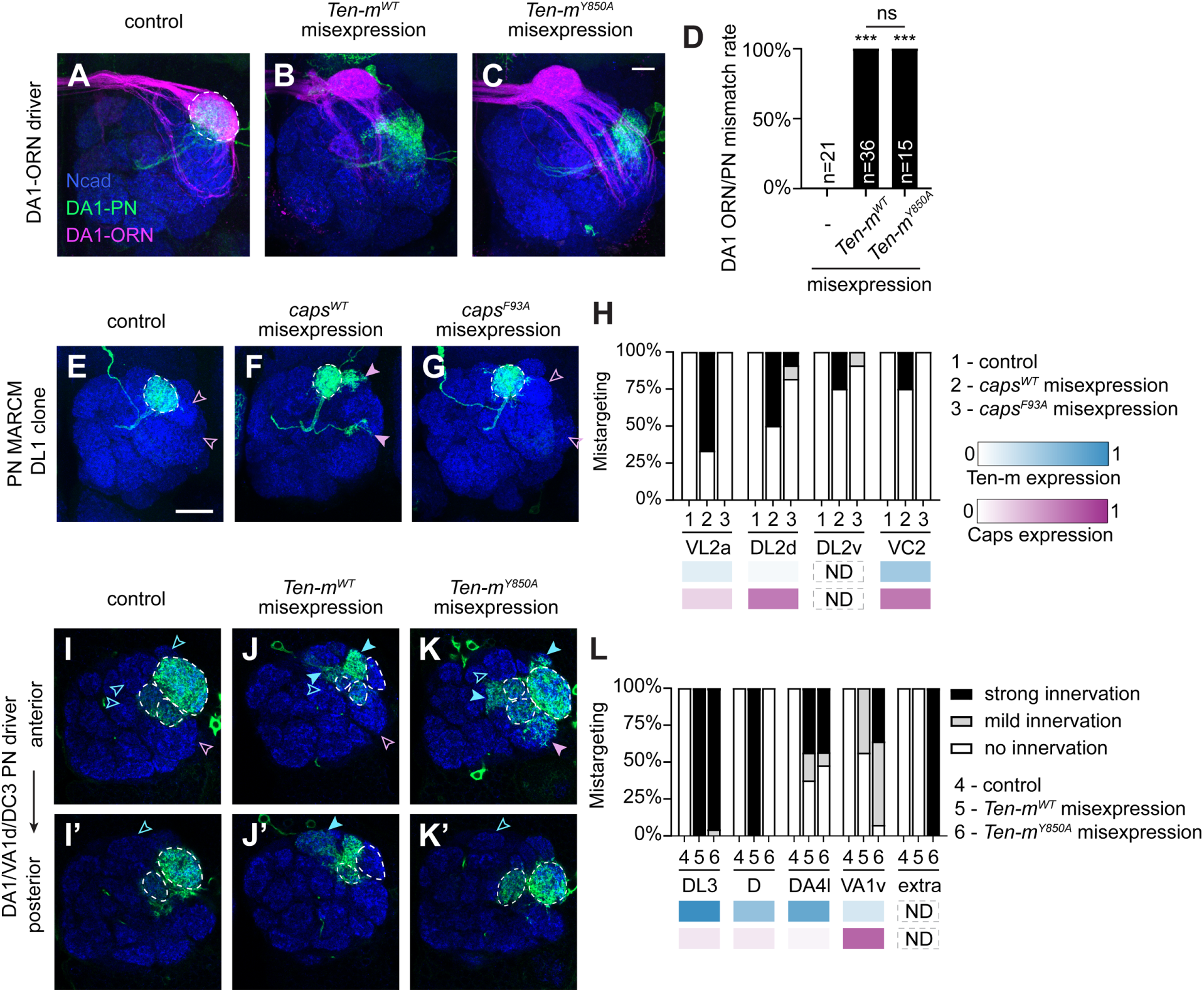
Gain-of-function analysis of Ten-m and Caps in ORN–PN matching and dendrite segregation, related to Figure 5. (A–C) Confocal images showing DA1-ORN axons and cognate DA1-PNs for control (A), *Ten-m^WT^*- misexpression in DA1-ORNs (B), and *Ten-m^Y850A^*-misexpression in DA1-ORNs (C). Instead of matching with DA1-PNs, misexpression of both *Ten-m^WT^* and *Ten-m^Y850A^* causes DA1-ORNs to mistarget to the DL3 glomeruli^13^. Scale bar, 10 μm. (D) Quantification of mistargeting penetrance. Statistical differences between conditions were assessed using 2×2 Fisher’s Exact Test followed by Benjamini-Hochberg (FDR) correction for all pairwise comparisons. ns (not significant); ***, p < 0.001. (E–G) Confocal images showing DL1-PN MARCM clones for control (E), *caps^WT^* misexpression (F), and *caps^F93A^*misexpression (G). Open/filled pink arrowheads indicate normally uninnervated and ectopically innervated Ten-m-low regions, respectively. Misexpression of *caps^WT^* causes DL1-PN dendrites to invade Ten-m-low regions, whereas *caps^F93A^* misexpression shows minimal effect. Scale bar: 20 μm. (H) Quantification of DL1-PN mistargeting penetrance for genotypes shown in E–G, with corresponding protein expression profiles at 48 h APF (n = 9, 12, 11 antennal lobes, respectively). (I–K’) Confocal images showing Mz19-PN dendrites (*Mz19-GAL4*) for control (I, I’), *Ten-m^WT^* misexpression (J, J’), and *Ten-m^Y850A^* misexpression (K, K’). Open/filled arrowheads indicate normally uninnervated and ectopically innervated Caps-low (blue) or Caps-high (pink) regions, respectively. Control Mz19-PN dendrites target DA1, VA1d, DC3 glomeruli. *Ten-m^WT^*misexpression causes Mz19 dendrites to mistarget to Caps-low DL3, D, DA4l regions (filled blue arrowheads in J, J’). *Ten-m^Y850A^* misexpression causes mistargeting to Caps-low DL3, the formation of an extra, ectopic glomerulus (not present in control brains; filled blue arrowheads in K), as well as Caps-high VA1v (filled pink arrowhead in K). (L) Quantification of Mz19-PN mistargeting penetrance for genotypes shown in I–K’, with corresponding protein expression profiles at 48 h APF (n = 20, 16, 23 antennal lobes, respectively).

**Table S1.**
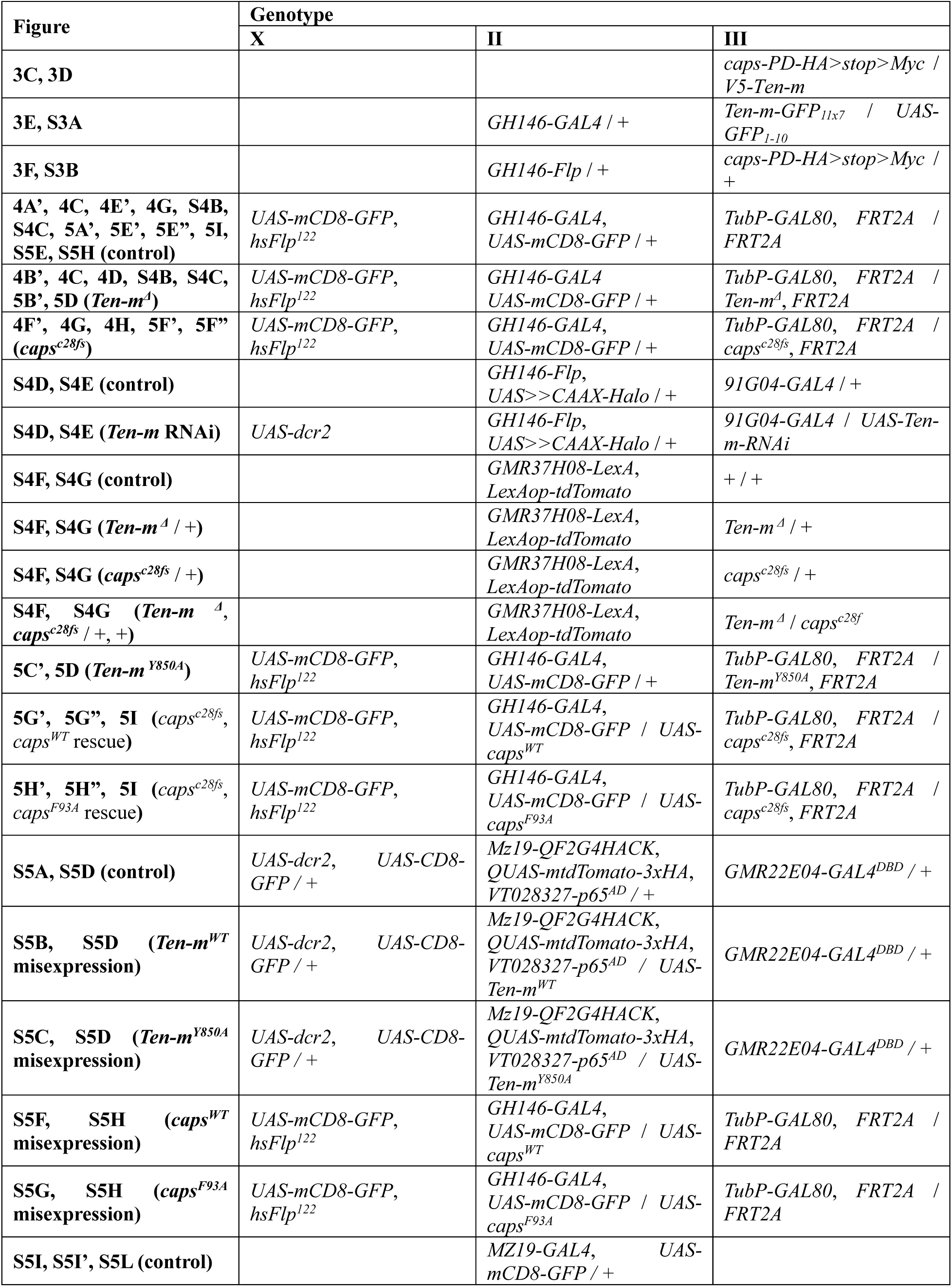
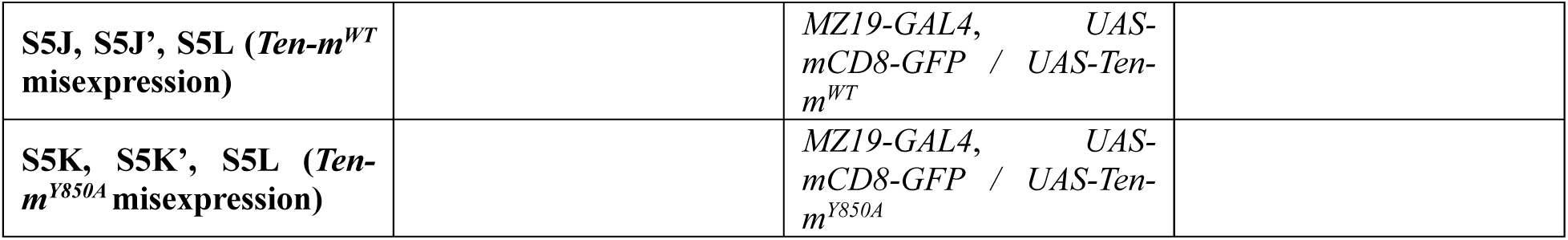
Complete genotypes of each experiment, related to STAR Methods.

## STAR Methods

### Key Resources Table

| REAGENT or RESOURCE | SOURCE | IDENTIFIER |
| --- | --- | --- |
| **Antibodies** |  |  |
| rat anti-DNcad | Developmental Studies Hybridoma Bank | Cat# DN-Ex #8; RRID: AB_528121 |
| chicken anti-GFP | Aves Labs | Cat# GFP-1020; RRID: AB_10000240 |
| rabbit anti-DsRed | Takara Bio | Cat# 632496; RRID: AB_10013483 |
| Mouse anti-GFP (clone GFP-20) | Sigma-Aldrich | Cat# G6539; RRID: AB_259941 |
| mouse anti-V5 | Thermo Fisher Scientific | Cat# R960-25; RRID: AB_2556564 |
| rabbit anti-HA | Cell Signaling | Cat# 3724; RRID: AB_1549585 |
| rabbit anti-Myc | Cell Signaling | Cat# 2278S; RRID: AB_490778 |
| rabbit anti-Halo | Promega | Cat# G9281 |
| Alexa Fluor 594-conjugated anti-HRP | Jackson ImmunoResearch | Cat# 123-585-021 |
| Hoechst 33342 | Thermo Fisher Scientific | Cat# H3570 |
| anti-His-iFlour 488 | GenScript | Cat# A01800 |
| **Chemicals, peptides, and recombinant proteins** |  |  |
| Schneider's *Drosophila* medium | Thermo Fisher Scientific | Cat# 21720001 |
| Fetal Bovine Serum, heat inactivated | Thermo Fisher Scientific | Cat# A3840101 |
| Effectene transfection reagent | Qiagen | Cat# 301425 |
| **Critical commercial assays** |  |  |
| Q5 hot-start high-fidelity DNA polymerase | New England Biolabs | Cat# M0494S |
| NEBuilder HiFi DNA assembly master mix | New England Biolabs | Cat# E2621L |
| **Deposited data** |  |  |
| scRNA-seq dataset | This study | GEO: GSE318611 |
| *Drosophila*-PN-scRNAseq-v1.0.0 | This study | DOI: 10.5281/zenodo.20941706 |
| **Experimental models: Organisms/strains** |  |  |
| *D. melanogaster*: APEX2-V5-Ten-m | Xu et al.<sup>13</sup> | N/A |
| *D. melanogaster*: GH146-GAL4 | Stocker et al.<sup>45</sup> | BDSC: 30026 |
| *D. melanogaster*: GH146-Flp | Hong et al.<sup>9</sup> | N/A |
| *D. melanogaster*: Mz19-GAL4 | Ito et al.<sup>46</sup> | BDSC: 34497 |
| *D. melanogaster*: 91G04-GAL4 | Jenett et al.<sup>47</sup> | N/A |
| *D. melanogaster*: GMR22E04-GAL4<sup>DBD</sup> | Jenett et al.<sup>47</sup> | BDSC: 69199 |
| *D. melanogaster*: VT028327-p65<sup>AD</sup> | Tirian et al.<sup>48</sup> | BDSC: 73064 |
| *D. melanogaster*: GMR37H08-LexA | Pfeiffer et al.<sup>49</sup> | BDSC: 52766 |
| *D. melanogaster*: D42-GAL4 | Sanyal et al.<sup>50</sup> | N/A |
| *D. melanogaster*: VT033006-GAL4 | Tirian et al.<sup>48</sup> | BDSC: 73333 |
| *D. melanogaster*: UAS-mCD8-GFP | Lee et al.<sup>29</sup> | DGRC: 108068 |
| *D. melanogaster*: UAS-dcr2 | Dietzl et al.<sup>51</sup> | N/A |
| *D. melanogaster*: QUAS-mtdTomato-3xHA | Potter et al.<sup>52</sup> | BDSC: 30004 |
| *D. melanogaster*: LexAop-tdTomato | Poe et al.<sup>53</sup> | BDSC: 77138 |
| *D. melanogaster*: hsFlp<sup>122</sup> | Golic et al.<sup>54</sup> | N/A |
| *D. melanogaster: TubP-GAL80,FRT2A* | Lee et al.<sup>29</sup> | N/A |
| *D. melanogaster: FRT2A* | Chou et al.<sup>30</sup> | N/A |
| *D. melanogaster: UAS-Ten-m-RNAi* | VDRC | VDRC: v51173 |
| *D. melanogaster: V5-Ten-m* | Xu et al.<sup>13</sup> | N/A |
| *D. melanogaster: Ten-m-GFP<sub>11x7</sub>* | Kamiyama et al.<sup>27</sup> | N/A |
| *D. melanogaster: UAS-GFP<sub>1-10</sub>* | Kamiyama et al.<sup>55</sup> | N/A |
| *D. melanogaster: UAS-FRT10-stop-FRT10-3xHalo7-CAAX* | Wong et al.<sup>8</sup> | N/A |
| *D. melanogaster: caps<sup>c28fs</sup>* | Shinza-Kameda et al.<sup>56</sup> | N/A |
| *D. melanogaster: nos-Cas9* | Ren et al.<sup>57</sup> |  |
| *D. melanogaster: vas-Cas9* | Gratz et al.<sup>58</sup> |  |
| *D. melanogaster: Cre* | BDSC | BDSC:1501 |
| *D. melanogaster: caps-PD-HA>stop>Myc* | This study | N/A |
| *D. melanogaster: Ten-m<sup>Δ</sup>,FRT2A* | This study | N/A |
| *D. melanogaster: Ten-m<sup>Y850A</sup>,FRT2A* | This study | N/A |
| *D. melanogaster: Mz19-QF2<sup>G4HACK</sup>* | This study | N/A |
| *D. melanogaster: UAS-Ten-m<sup>WT</sup>* | This study | N/A |
| *D. melanogaster: UAS-Ten-m<sup>Y850A</sup>* | This study | N/A |
| *D. melanogaster: UAS-caps<sup>WT</sup>* | This study | N/A |
| *D. melanogaster: UAS-caps<sup>F93A</sup>* | This study | N/A |
| *D. melanogaster: S2 cells* | Thermo Fisher Scientific | Cat# R69007 |
| **Recombinant DNA** |  |  |
| *pU6-caps-FRT-3xHA-stop-FRT-3xMyc-mCherry donor* | This study | N/A |
| *pU6-Ten-m<sup>Δ</sup>-DsRed donor* | This study | N/A |
| *pU6-Ten-m<sup>Y850A</sup>-DsRed donor* | This study | N/A |
| *pU6-Ten-m<sup>Y850A</sup> gRNA* | This study | N/A |
| tagged *UAS-Ten-m* and *-Ten-m<sup>ECD</sup>* constructs | This study |  |
| tagged *UAS-caps* and *-caps<sup>ECD</sup>* constructs | This study |  |
| **Software and algorithms** |  |  |
| Zen | Carl Zeiss | RRID: SCR_013672 |
| ImageJ | National Institutes of Health | RRID: SCR_003070 |
| Illustrator | Adobe | RRID: SCR_010279 |
| STAR (2.7.10a) | Dobin et al.<sup>59</sup> | RRID: SCR_015899 |
| Iterative Clustering for Identifying Markers (ICIM) | Li et al.<sup>43</sup> | N/A |
| Scanpy (1.11.4) | Wolf et al.<sup>60</sup> | RRID:SCR_018139 |
| ColabFold (v1.5.5) | Mirdita et al.<sup>23</sup> | RRID:SCR_025453 |
| MODELLER (v 10.6) | Webb et al.<sup>61</sup> | N/A |
| PyMOL (v. 3.1.6) | Schrödinger, LLC | RRID:SCR_000305 |
| UCSF Chimera (v. 1.11) | Pettersen et al.<sup>62</sup> | RRID:SCR_004097 |
| GraphPad Prism | GraphPad Software | RRID:SCR_002798 |
| SnapGene | Insightful Science | RRID:SCR_015053 |

### Experimental Model and Participant Details

#### *Drosophila* stocks and genotypes

*Drosophila melanogaster* were reared on standard cornmeal molasses agar medium at 25°C under a 12-h light/12-h dark cycle. For experiments involving MARCM or heat-shock induction, specific temperature shifts were applied as detailed below. Complete genotypes for flies used in each experiment are described in Table S1.

### Method Details

#### Generation of *Ten-m^Δ^* and *Ten-m^Y850A^* flies

*Ten-m^Δ^* was generated by CRISPR/Cas9-mediated replacement of exons 3–6 with a selection cassette. A self-linearizing donor construct was designed in which a *loxP-DsRed-loxP* cassette flanked by homology arms (HR1 upstream of exon 3, HR2 downstream of exon 6) was itself flanked by the same gRNA target sequence that cleaves the genomic locus. A pU6-driven gRNA cassette on the same vector directs Cas9 to simultaneously linearize the donor and cut the genomic target, facilitating homology-directed repair^38^. The construct was injected into *vas-Cas9*;*FRT2A*^58,63^ embryos. Integrants were identified by DsRed fluorescence. The deletion removes Ten-m’s sole transmembrane domain and introduces a frameshift C-terminal to the deletion. Correct deletion was verified by Splinkerette PCR^64^ followed by Sanger sequencing. *Ten-m^Y850A^* was generated by CRISPR/Cas9-mediated knock-in using a self-linearizing donor strategy. Two gRNAs were used: one encoded on the donor plasmid targeting sequences downstream of the Y850 codon, and a second on a separate pU6 plasmid targeting upstream of Y850. The donor construct contained a 5’ homology arm immediately upstream of Y850, followed by the replacement sequence harboring the Y850A mutation (TAC→GCA) and the remainder of exon 9, a *loxP-DsRed-loxP* selection cassette inserted within the intron downstream of exon 9, and a 3’ homology arm beginning from the same intron. Silent mutations were introduced at the PAM sites to prevent re-cutting. The two plasmids were co-injected into *vas-Cas9*;*FRT2A* embryos. Integrants were identified by DsRed fluorescence, and the selection cassette was removed by crossing to a line carrying balancer expressing Cre (Bloomington *Drosophila* Stock Center, RRID:BDSC 1501). Correct knock-in was verified by Sanger sequencing.

#### Generation of conditional tag line

The *caps-PD-HA>stop>Myc* conditional allele was generated using a similar CRISPR/Cas9-mediated knock-in strategy as described in^28,65^. In brief, a donor construct containing *HR1-FRT-3xHA-stop-FRT- 3xMyc-stop-loxP-mCherry-loxP-HR2* and a pU6-driven gRNA cassette was injected into *nos-Cas9* embryos^57^. Integrants were identified by mCherry fluorescence. The *loxP-mCherry-loxP* cassette was removed by Cre (BDSC 1501). In the default state, Caps protein is tagged with 3xHA; upon *FLP*-mediated recombination (e.g., via *GH146-FLP*), the HA-stop cassette was excised and Caps is tagged with 3xMyc, enabling PN-specific detection.

#### Generation of UAS constructs and transgenic flies

Tagged *UAS-Ten-m* and *-ECDs* were generated from multiple gBlocks synthesized by Twist Biosciences and cloned into a 10× *pUASt-attB* vector using Gibson Assembly. Tagged *UAS-caps* and -ECDs were synthesized and cloned by Twist Biosciences into a 10× *pUASt-attB* vector. All constructs were validated by full-length plasmid sequencing. The *UAS-Ten-m* constructs were injected in house into embryos bearing the attP24 landing site. The UAS-caps constructs were made through BestGene Inc and injected into embryos bearing the VK37 site. G0 flies were crossed to a *w⁻* balancer, and all *w⁺* progeny were individually balanced.

### Single-cell transcriptome acquisition and analysis

#### Single-cell RNA sequencing procedure

Single-cell RNA sequencing was performed following previously described protocol^20,43^. Briefly, *Drosophila* brains with *mCD8-GFP* labeled cells using *VT033006-GAL4* drivers were dissected at 12–18 h APF. Single-cell suspension was prepared and GFP positive cells were sorted using Fluorescence Activated Cell Sorting (FACS) into individual wells of 384-well plates containing lysis buffer using SH800 (Sony Biotechnology). Full-length poly(A)-tailed RNA was reverse-transcribed and amplified by PCR following the SMART-seq2 protocol^66^. cDNA was digested using lambda exonuclease (New England Biolabs) and then amplified for 25 cycles. Sequencing libraries were prepared from amplified cDNA, pooled, and quantified using BioAnalyser (Agilent). Sequencing was performed using the Novaseq 6000 Sequencing system (Illumina) with 100 paired-end reads and 2 × 12 bp index reads.

#### Sequence processing and analysis

Reads were aligned to the *Drosophila melanogaster* genome (r6.10) using STAR (2.7.10a)^59^. Gene counts were produced using STAR quantMode GeneCounts. Raw count tables from 384-well plates were subjected to a multi-stage filtering process: (1) exclusion of ERCC spike-in outliers, (2) removal of low-quality cells with library sizes < 100,000 counts or mitochondrial gene content > 5%, and (3) selection of cells expressing established neuronal markers to ensure population purity. The filtered count matrices were normalized to counts per million (CPM) and log-transformed. To facilitate cross-stage comparative analysis, the 12 h APF dataset was integrated with previously published PN transcriptomes (GSE161228; 0 h, 24 h, 48 h APF, and adult)^20^. Feature selection was performed using the Iterative Clustering for Identifying Markers (ICIM) algorithm^43^, specifically targeting genes critical for neuronal identity, such as transcription factors (TFs) and cell-surface molecules (CSMs). For visualization, high-dimensional data were embedded into two-dimensional space using t-Distributed Stochastic Neighbor Embedding (t-SNE). Initial graph-based partitioning of the single-cell transcriptomes was performed using the Leiden community detection algorithm (as implemented in Scanpy). To ensure the robustness of our cell-type clusters, we further employed HDBSCAN (Hierarchical Density-Based Spatial Clustering of Applications with Noise) to identify stable clusters within the t-SNE embedding space. To assign biological identities, we calculated the cosine similarity between the mean expression profiles of 12 h APF clusters and annotated PN types at 24 h APF. Final cell-type matching was refined through manual annotation using lineage-specific markers.

All data analysis was performed in Python using Scanpy, NumPy, SciPy, Pandas, scikit-learn, and the hdbscan package. Specific single-cell RNA-seq computational workflows and matching algorithms were implemented using custom modules and established pipelines as previously described^20,43,44^. Sequencing reads and preprocessed sequence data are available in the NCBI Gene Expression Omnibus (GSE318611). Custom analysis code is available at https://github.com/phagocyt/FlyPN.

#### Anti-correlation analysis of cell-surface molecules

To identify molecular pairs with complementary or antagonistic expression patterns, we performed a systematic correlation analysis focusing on cell-surface molecules (CSMs) across projection neuron (PN) subtypes at 12 h APF. The candidate CSM list was obtained from^67^. Data were first aggregated into subtype-level profiles by calculating the mean log-transformed CPM (expression intensity) and the expression fraction (percentage of cells with detected transcripts) for each PN type. We applied a strict filter cutoff, retaining only CSMs with a mean expression > 3 log_2_(CPM+1) and an expression fraction > 50% in at least one subtype. For target gene (e.g., *caps*), we computed the Pearson correlation coefficient (*r*) against all filtered CSMs. An integrated correlation score was derived by averaging the coefficients from the intensity (*r_cpm_*) and breadth (*r_frac_*) matrices:

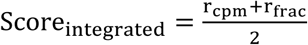

CSMs were sorted in ascending order by this score. Genes exhibiting the strongest anti-correlation (scores closest to -1) were identified as top candidates for structural analysis.

### Structural prediction and modeling

#### Colabfold predictions

Interactions between the Caps ectodomain (ECD) and the ECDs of the top 10 anti-correlated candidates were predicted using ColabFold (v1.5.5), an accelerated implementation of AlphaFold2^23^. Structure predictions were performed using the alphafold2_multimer_v3 model weights, 5 models with 20 recycles, templates enabled, and no AMBER relaxation. The Multiple Sequence Alignments (MSAs) are acquired through MMseqs2 MSA server. Complexes were ranked based on the average inter-chain predicted Template Modeling (ipTM) scores of five models. Predictions were run on 80 GB H100 NVIDIA GPUs.

#### Homology modeling

We used MODELLER version 10.6 for homology modeling of the Ten-m–Caps complex. As known inputs, we provided MODELLER with our *C. elegans* TEN-1–LRON-11 structure from our companion manuscript (Li et al., companion manuscript), the crystal structure of Ten-m^37^ and the Alphafold models of Ten-m and Caps separately to aid with missing loops and domains in the experimentally determined structures. The resulting model and the Colabfold models were used for mutational design to break the Ten-m–Caps complexes.

#### Extracellular interactome assay (ECIA)

Pairwise protein interaction assay was applied as described previously^24^. Briefly, wild-type and mutant constructs covering the extracellular domain of Ten-m, including a WT construct with a splice insertion in the TTR domain (SIFWNYFNA) found in isoform E, were tagged with an Fc tag, and secreted to be used as bait on Protein A-coated 96-well plates. Alkaline phosphatase (AP)-tagged Caps ectodomains were secreted and used as prey. Interaction was detected by AP activity using absorbance at 650 nm from the AP substrate BluePhos (KPL, 50-88-02). Fc-tagged Ten-m ECD (WT, Y850A, and +SS isoform E splice variant) and AP-tagged Caps ECD (WT, F93A) constructs were generated by the Araç laboratory as described^24,25^. *Drosophila* Rst ECD-Fc and Rst ECD-AP were used as negative controls^25^. Expression levels for each mutant series and their corresponding wild-type versions were quantified via Western blot analysis. This was performed using an anti-His-Tag Antibody coupled with iFlour 488 (Genscript, A01800, 1:5000 dilution), targeting the 6xHis tags engineered at the C-termini of all constructs. This blotting also confirmed the successful expression and secretion of the mutant ectodomains. Fluorescent signal quantification was performed using a Biorad Chemidoc system and ImageLab version 6 (Biorad). Normalization of protein quantities across samples was achieved by diluting higher-expressing variants with fresh Schneider’s culture media to match the expression levels of lower-expressing variants.

### Transfection and immunostaining of *Drosophila* S2 cells

#### Cell-based binding assay

*Drosophila* S2 cells (Thermo Fisher Scientific) were cultured in Schneider’s medium supplemented with 10% heat-inactivated FBS (Thermo Fisher Scientific) at 25°C. To produce soluble extracellular domain (ECD) proteins, cells in 6-well plates were transfected with *Actin-GAL4* and V5-tagged ECD constructs (*Ten-m^ECD^-V5*, *caps^ECD^-V5*) using Effectene (Qiagen). Conditioned medium was collected 3 days post-transfection. Separately, S2 cells on poly-D-lysine-coated coverslips (German coverslips) in 24-well plates were transfected with *Actin-GAL4* and full-length receptor constructs: *Halo- Ten-m^WT^*, *Halo-Ten-m^Y850A^*, *caps^WT^-HA* or *caps^F93A^-HA*. Two days post-transfection, conditioned medium containing the appropriate soluble ECD was applied overnight at room temperature. Cells were washed with PBS, fixed in 4% PFA for 15 min, permeabilized, and stained with anti-V5 (1:200; Thermo Fisher Scientific) to detect ECD binding and anti-HA (1:300; Cell Signaling), or anti-Halo (1:500; Promega) to identify receptor-expressing cells. Secondary antibodies (1:500; Jackson ImmunoResearch) and DAPI were applied, and coverslips were mounted in SlowFade (Thermo Fisher Scientific). Images were acquired on a Zeiss LSM 900 with a 20× or 63× objective.

#### Live surface staining

To verify that binding-deficient mutations did not impair membrane trafficking (Figures S2D–S2G), transfected S2 cells on coverslips were stained prior to fixation under non-permeabilized conditions. Medium was gradually replaced with pre-chilled Schneider’s medium, and cells were incubated with anti-HA (1:30; Cell Signaling, C29F4) or anti-V5 (1:20; Thermo Fisher Scientific, R960-25) diluted in cold medium for 30 min at 4°C. Cells were washed twice with cold PBS, fixed in 4% PFA for 10 min, and incubated with Alexa Fluor-conjugated secondary antibodies (1:500; Jackson ImmunoResearch) and Hoechst 33342 (1:1000) for 1 h at room temperature without permeabilization.

#### Immunostaining of fly brains

Fly brains were dissected according to a previously published protocol^68^. In brief, brains were dissected in PBS, transferred to a tube containing 4% paraformaldehyde in PBST (0.3% Triton X-100), and fixed for 20 min while nutating at RT. Following fixation, brains were washed 3 times for 20 min in PBST and blocked for at least 30 min in PBST + 5% normal donkey serum. The following antibodies were used: rat anti-Ncad (Developmental Studies Hybridoma Bank; 1:40), chicken anti-GFP (Aves Labs; 1:1000), rabbit anti-HA (Cell Signaling Technologies; 1:300), rabbit anti-MYC (1:300, 2278S, Cell Signaling), mouse anti-V5 (Thermo Fisher Scientific; 1:200), rabbit anti-dsRed (Takara Bio; 1:1000), and incubated with brains in block buffer overnight at 4°C while nutating. Brains were subsequently washed three times for 20 min in PBST and incubated in secondary antibodies (Alexa Fluor 488; Alexa Fluor Cy3; Alexa Fluor 647; 1:500) overnight at 4°C while nutating. Brains were again washed three times for 20 min in PBST, transferred to SlowFade antifade reagent (Thermo Fisher Scientific) and stored at 4°C prior to mounting.

#### Neuromuscular junction analysis

To confirm protein stability and axonal transport of Ten-m and Caps variants *in vivo* (Figures S2H–S2K’’), *UAS-Halo-Ten-m^WT^*, *UAS-Halo-Ten-m^Y850A^*, *UAS-caps^WT^-HA* or *UAS-caps^F93A^-HA* were expressed in motor neurons using *D42-GAL4*. Wandering L3 larvae were dissected and processed for immunofluorescence as previously described^69^. In brief, wandering L3 larvae were dissected in PBS and fixed in 4% PFA for 20 min. Larval fillets were washed in PBS with 0.3% Triton X-100, blocked in 5% normal donkey serum, and incubated with anti-Halo (1:500; Promega) or anti-HA (1:300; Cell Signaling) together with Alexa Fluor 594-conjugated anti-HRP (1:300; Jackson ImmunoResearch) overnight at 4°C. Secondary antibodies (1:500; Jackson ImmunoResearch) were applied for 2 h at room temperature.

#### Image acquisition and processing

Images were obtained on a Zeiss LSM900 laser-scanning confocal microscope (Carl Zeiss) using either a 40× or 63× oil immersion objective. 8-bit z-stacks were acquired at 1 μm intervals at a resolution of 1024 x 512. Brightness and contrast adjustments as well as image cropping were done using Photoshop (Adobe) or FIJI/Image J.

#### MARCM-based clonal analyses

Clonal analyses using mosaic analysis with a repressible cell marker (MARCM) have been previously described^70^. Each fly contains a *hsFLP^122^* recombinase, *GH146-GAL4* (PN *GAL4*), *TubP-GAL80*, *UAS- mCD8-GFP*, the *FRT2A*, and either wild-type or a mutant allele distal to the FRT site. To generate lateral PN (lPN) neuroblast clones and DL1 single-cell clones, flies were heat shocked for 1 hour at 37°C at 0–24 h after larval hatching. To generate single-neuron clones with projections to the VC1, DC2, or VA7m glomerulus, flies were heat shocked for 1 hour at 37°C at 24–48 h after larval hatching. To generate single-neuron clones with projections to the DA2 or DM5 glomerulus, flies were heat shocked for 1 hour at 37°C at 96–120 h after larval hatching. For DL1 single-cell misexpression analysis, flies were raised at 29°C after heat shock to increase expression level and overcome GAL80 perdurance. Brains were dissected and processed as described above.

### Quantification and Statistical Analysis

#### S2 cells fluorescence intensity analysis

ECD binding was quantified using an automated ImageJ macro. Confocal z-stacks taken by a 20× objective were max-projected and split into full-length receptor (FL), ECD, and DAPI channels. Transfected cells (FL-positive) were segmented by Otsu thresholding on the receptor channel. Untransfected negative control cells were identified by Otsu thresholding on the DAPI channel, subtracting the FL-positive mask to exclude transfected cells. Nuclear ROIs were dilated by ∼1.25 µm to approximate cell body boundaries, and regions overlapping with FL-positive cells were excluded. ROIs were filtered by area (7.8–23.4 µm²) and circularity (>0.9) to select single, round cells. Mean ECD fluorescence intensity was measured within FL-positive and negative control ROIs on the ECD channel. Statistical comparisons were performed using repeated measures two-way ANOVA followed by Šídák’s multiple comparisons test (Figures 2K, 2L) or paired *t*-test (Figures S2M and S2O).

#### Expression pattern profiling

For quantification of Ten-m and Caps protein distribution at 18 h APF (Figure 3D), a 10-pixel-wide straight line was drawn along the dorsomedial–ventrolateral (DM–VL) axis on the anterior side of the antennal lobe in Fiji/ImageJ. Fluorescence intensity values for the Ten-m and Caps channels were extracted, min-max normalized, and averaged across antennal lobes (n = 12). For quantification of PN-specific protein expression at 48 h APF (Figures 3E and 3F), individual glomeruli were manually segmented as ROIs. Mean fluorescence intensity of the Ten-m or Caps channel within each ROI was normalized to the Ncad channel to correct for depth-dependent signal attenuation (n = 5 antennal lobes).

#### Dendrite Mistargeting Scoring

Dendrite targeting was scored as a categorical variable (mistargeted or correctly targeted) by visual inspection of confocal z-stacks. A neuron was scored as mistargeted if dendrite branches extended into glomeruli outside the expected target. Scoring was performed blind to genotype where possible (Figures 4C, 4D, 4G, 4H, 5D, 5I, S4E, S4G, S5D, S5H, S5L).

#### Statistical analysis

All statistical tests were performed using GraphPad Prism. For cell-based binding assays, repeated measures two-way ANOVA followed by Šídák’s multiple comparisons test (Figures 2K and 2L) or paired t-test (Figures S2M and S2O) was used. For all dendrite mistargeting and ORN mismatching experiments, penetrance was compared using two-sided Fisher’s exact test on 2×2 contingency tables. For experiments involving multiple comparisons, p-values were adjusted using the Benjamini-Hochberg False Discovery Rate (FDR) procedure (Figures 4C, 4G, 5I, S4E, S4G, S5D, S5H, S5L). Correlation between *Ten-m^Y850A^* and *Ten-m^Δ^*glomerular innervation frequencies was assessed using Pearson’s correlation coefficient (Figure 5D). Significance levels: *p < 0.05, **p < 0.01, ***p < 0.001; ns, not significant. Data are presented as mean ± SEM unless otherwise noted.

